# CaMKII–LUZP1 signaling couples cytoskeletal acetylation and autophagy to drive neuronal plasticity

**DOI:** 10.64898/2026.07.03.736264

**Authors:** Chao-Yuan Tsai, Kazuki Kuroda, Yuichiro Oka, Manabu Taniguchi, Hideshi Yagi, Makoto Sato

## Abstract

Cytoskeletal acetylation and autophagy are fundamental drivers of neuronal plasticity, yet how these pathways are coordinated across subcellular compartments remains unknown. Here, we identify LUZP1 as a signaling organizer that couples cytoskeletal acetylation to autophagy in hippocampal neurons. LUZP1 deficiency impaired neurite outgrowth and dendritic spine maturation while promoting ciliary elongation; these phenotypes were partially mirrored in neuron-specific *Luzp1* knockout mice, which also showed altered locomotor behavior. Mechanistically, LUZP1 promoted neurite extension by enhancing ATAT1-dependent α-tubulin acetylation while driving spine maturation and limiting ciliary growth by restraining HDAC6-dependent cortactin (CTTN) deacetylation. An acetylation-mimetic CTTN mutant rescued both spine and ciliary defects caused by LUZP1 deficiency. In parallel, blocking autophagy-dependent OFD1 degradation attenuated ciliary elongation, linking CTTN deacetylation to increased autophagy under LUZP1-deficient conditions. Finally, activated CaMKIIα associated with LUZP1 and selectively enhanced its interaction with HDAC6 and CTTN, coupling neuronal activity to cytoskeletal remodeling. Together, these findings identify a CaMKIIα–LUZP1 pathway that integrates cytoskeletal acetylation with autophagy to coordinate neuronal morphogenesis and ciliary homeostasis.

**Highlights:**

- LUZP1 couples cytoskeletal acetylation to autophagy in hippocampal neurons
- LUZP1 promotes neurite extension through ATAT1-dependent α-tubulin acetylation
- LUZP1 restrains HDAC6–CTTN signaling to drive spine maturation and limit ciliary growth
- CaMKIIα selectively strengthens the LUZP1–HDAC6–CTTN pathway

## Introduction

The neuronal cytoskeleton provides a dynamic framework for brain development and neural function across the lifespan. Microtubules and actin filaments are core determinants of neuronal polarization, neurite extension, and synaptic plasticity^1,2^. Post-translational modifications of cytoskeletal proteins, particularly acetylation^3^, have emerged as a key mechanism controlling neuronal development and neural function^4,5^. α-Tubulin acetylation, governed by acetyltransferases such as α-tubulin acetyltransferase 1 (ATAT1) and deacetylases such as histone deacetylase 6 (HDAC6), regulates microtubule stability and dynamics^6–10^. In parallel, acetylation of cortactin (CTTN) controls its interactions with F-actin and associated cytoskeletal factors, thereby shaping actin-dependent remodeling^11–14^. Disruption of acetylation homeostasis impairs core neurodevelopmental processes, including neuronal migration^15,16^, axon guidance^17–19^, dendritic arborization^16,20^, and synapse formation^11,20,21^, ultimately compromising neural circuit assembly and refinement^7,16,20,22,23^.

Autophagy has likewise emerged as a central regulator of neuronal development and plasticity^24^. Beyond its canonical role in protein and organelle turnover, autophagy contributes to the dynamic remodeling of specialized neuronal compartments, including axons, growth cones, dendritic spines, and primary cilia^25–28^. Because autophagic activity is tightly linked to acetylation-dependent cytoskeletal regulation^29^, these pathways are well positioned to act together in shaping neuronal architecture. However, the mechanisms that couple cytoskeletal acetylation to autophagy across distinct neuronal compartments remain poorly defined.

Leucine zipper protein 1 (LUZP1) has emerged as an important regulator of cytoskeletal organization and neuronal development^30,31^. This brain-enriched protein, expressed in cortical and hippocampal neurons^32^, influences multiple features of cytoskeletal architecture. Genetic deficiency of LUZP1 has been linked to developmental disorders, including Townes–Brocks syndrome and 1p36 deletion syndrome, which are associated with defective ciliogenesis, abnormal cell migration, and impaired centrosome function^33–35^. At the mechanistic level, LUZP1 bundles F-actin and binds mechanically unfolded filamin A (FLNA) repeat-21, functioning as a tension-sensitive integrator of actin- and FLNA-dependent mechanotransduction^36^. LUZP1 also localizes to tight junctions, associates with microtubules, and sustains di-phosphorylated myosin light chain-driven actomyosin contractility by inhibiting myosin phosphatase, thereby promoting apical constriction during epithelial morphogenesis^37^. Together, these observations position LUZP1 as a candidate organizer of actin, microtubules, and tension-dependent signaling, raising the possibility that it also coordinates acetylation-dependent cytoskeletal remodeling in neurons.

Calcium/calmodulin-dependent protein kinase II (CaMKII) is a major activity-dependent kinase in neurons and a central regulator of long-term potentiation (LTP), dendritic spine remodeling, and synaptic plasticity^38^. In hippocampal neurons, CaMKII is activated downstream of N-methyl-D-aspartate receptor (NMDAR)-mediated Ca^2+^ influx and regulates F-actin organization by directly bundling actin or remodeling actin-associated structures within dendritic spines^39,40^. Activated CaMKII also translocates to dendritic microtubules, where it supports local synaptic plasticity^41^. In zebrafish, CaMKII is present in ciliated tissues and contributes to primary cilium stabilization^42^. Because both dendritic spine maturation and ciliogenesis require coordinated control of actin architecture and microtubule dynamics^2,43–45^, CaMKII is well positioned to couple neuronal activity to structural remodeling pathways. These observations raise the possibility that CaMKII converges with LUZP1-dependent mechanisms to coordinate compartment-specific remodeling during neuronal development and plasticity.

Despite emerging evidence linking LUZP1 to synaptic plasticity and brain function^46^, how it controls cytoskeletal remodeling remains unclear. Here, we identify LUZP1 as a signaling organizer that couples cytoskeletal acetylation to autophagy in hippocampal neurons. Through interactions with ATAT1, HDAC6, and CTTN, LUZP1 engages distinct pathways that govern neurite extension, dendritic spine maturation, and ciliary growth. We further show that CaMKIIα interacts with and phosphorylates LUZP1, selectively strengthening its association with CTTN and HDAC6, but not ATAT1, thereby linking neuronal activity to compartment-specific cytoskeletal remodeling. Together, these findings define a CaMKIIα–LUZP1 pathway that integrates cytoskeletal acetylation with autophagy to coordinate neuronal morphogenesis and ciliary homeostasis.

## Results

### LUZP1 is required for neurite extension and spine maturation in hippocampal neurons

To define the contribution of LUZP1 to neuronal morphogenesis, we first asked whether it is required for neurite growth and dendritic spine maturation. We generated two independent short hairpin RNAs (shRNAs) targeting mouse *Luzp1* mRNA. Immunoblot analysis showed that the LUZP1 shRNA constructs KD1 and KD2, which target nucleotide positions 888 and 2252, respectively, efficiently reduced exogenous mouse LUZP1 expression relative to scramble controls (Figure 1A). Immunofluorescence analysis confirmed reduced LUZP1 levels in the soma and dendrites of mouse hippocampal neurons transfected with the shRNA plasmids (Figure 1B).

**Figure 1.**
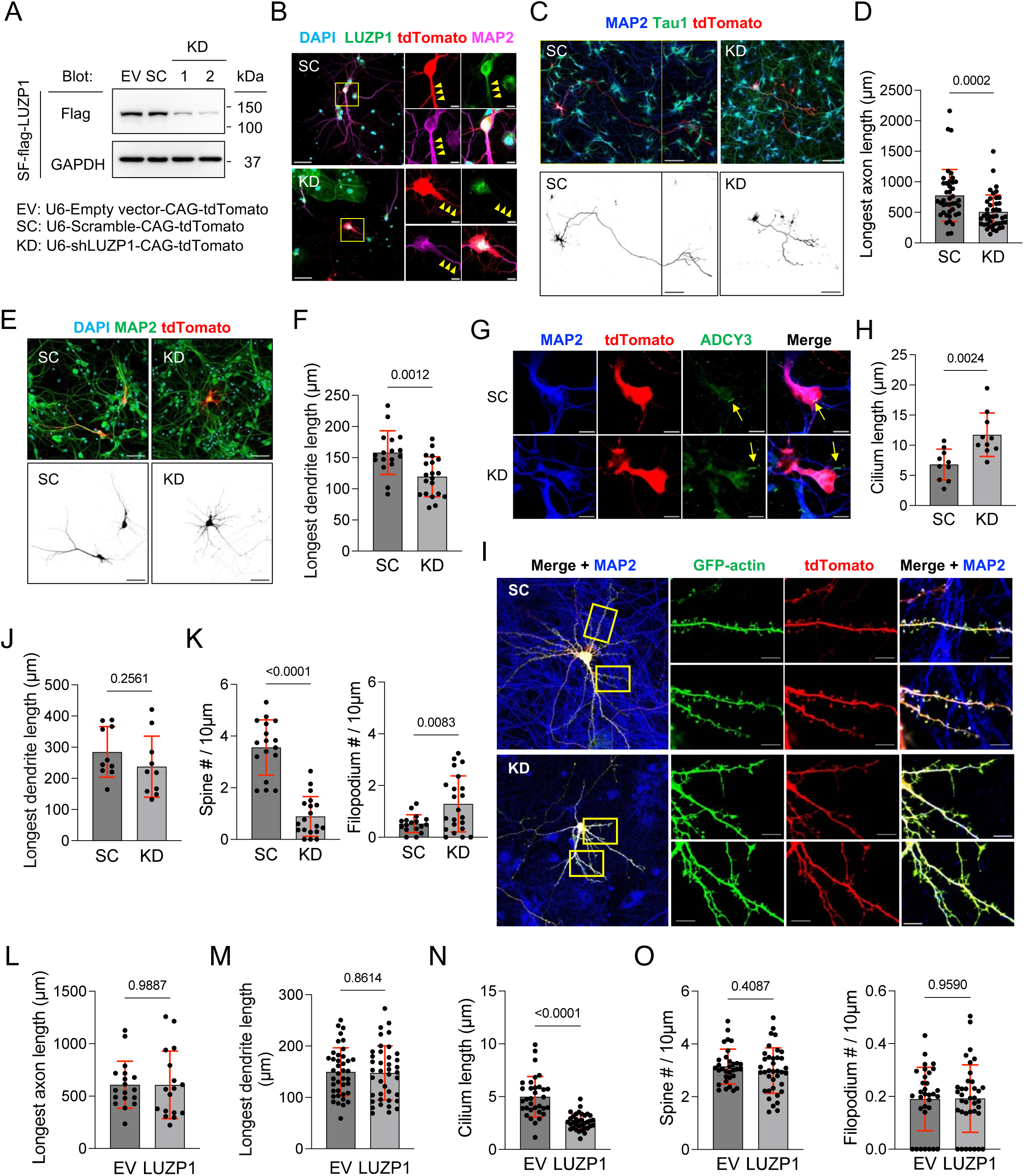
LUZP1 knockdown alters axon, dendrite, and ciliary growth, and spine maturation in cultured hippocampal neurons. (A) Validation of LUZP1 knockdown in HEK293T cells. pLKO.1-U6-shRNA-CAG-tdTomato shRNA vectors were cotransfected with either Strep-tag–Flag mouse LUZP1-expressing (SF-LUZP1) vectors or its empty vectors (EV), and LUZP1 expression was assessed using anti-Flag antibody. KD1 and KD2, independent targeting sequences; SC, scramble control. (B) LUZP1 immunostaining in hippocampal neurons transfected with SC or KD1 at DIV7 and fixed at DIV10. Yellow arrowheads indicate dendritic LUZP1. (C and D) Axon analysis in DIV5 hippocampal neurons knocked down at DIV3. Longest axon length was quantified. The images shown in the upper and lower panels were merged and tdTomato-only images, respectively. Images in SC-transfected hippocampal neurons were composed of two neighboring scanning fields. (E and F) Dendrite analysis in DIV10 hippocampal neurons knocked down at DIV7. Longest dendrite length was quantified. The images shown in the upper and lower panels were merged and tdTomato-only images, respectively. (G and H) Cilia analysis in DIV12 hippocampal neurons knocked down at DIV9. Cilium length was quantified. Yellow arrows indicate ADCY3^+^ cilia. (I) Dendrite and dendritic spine analysis in hippocampal neurons. Either SC or KD1 shRNA vectors were cotransfected with pCAG-mGFP-Actin vectors into hippocampal neurons at DIV15 and fixed at DIV18. (J-K) Quantification of dendrites and dendritic spines in DIV17-18 hippocampal neurons knocked down at DIV15. Longest dendrite length (J), and spine and filopodium densities (K) were quantified. Spines included thin, stubby, and mushroom morphologies. (L–O) Effects of LUZP1 overexpression on axon length (L, DIV3–5), dendrite length (M, DIV7–10), cilium length (N, DIV9–12), and spine density (O, DIV17–20). Scale bars, 50 μm and 10 μm (B, inset), 100 μm (C), 50 μm (E), and 10 μm (G and I). Each dot represents an individual neuron (D, F, H, J and L–N) or a dendritic branch segment from an individual neuron (2–7 neurons per condition; K and O). Data were from one represented experiment (L and M) and two independent experiments showed similar results. Data were pooled from two or three independent experiments (D, F, H, J, K, N, and O). Data are presented as mean ± SD. Two-tailed unpaired t-tests (D, F, H and J-L). Exact P values are shown.

We next performed stage-specific LUZP1 knockdown in primary hippocampal neurons (Figure 1C-1K). LUZP1 depletion at DIV3 and DIV7 reduced axonal and dendritic length at DIV5 and DIV10, respectively (Figures 1C–1F), whereas knockdown at DIV9 promoted ciliary elongation at DIV12 (Figures 1G and 1H). In addition, LUZP1 knockdown at DIV15 significantly reduced dendritic spine density at DIV18 without altering dendritic length (Figures 1I–1K). By contrast, LUZP1 overexpression reduced cilium length, but did not significantly affect axonal length, dendritic length, or dendritic spine density relative to control neurons (Figures 1L–1O and Figure S1). Together, these data identify LUZP1 as a stage-dependent regulator of neurite extension, spine maturation, and ciliary growth in hippocampal neurons.

### Neuron-specific loss of LUZP1 impairs locomotor behavior in mice

To test whether these phenotypes extend *in vivo*, we first generated *Luzp1*^tm1b^ reporter mice by inserting a β-galactosidase reporter cassette downstream of exon 3 of the *Luzp1* locus (Heterozygous, Figure 2A). We then generated neuron-specific *Luzp1* knockout (LUZP1-cKO) mice by crossing *Emx1*-Cre mice^47^ with *Luzp1*^tm1c^ (*Luzp1*^flox/flox^) mice (Figure 2A). X-gal staining and *in situ* hybridization showed that *Luzp1* is expressed predominantly in the hippocampus (Figure 2B and Figure S2A), and immunohistochemistry confirmed efficient loss of LUZP1 in LUZP1-cKO brains (Figure 2C). Immunoblot analysis further showed that LUZP1 was enriched in membrane fractions and was undetectable in LUZP1-deficient hippocampi (Figure S2B). In the open-field test, LUZP1-cKO mice displayed reduced locomotor activity and exploratory behavior, including fewer center entries, fewer rearing events, and reduced time spent in the center zone (Figures 2D–2G). Consistent with a role for LUZP1 in synaptic maturation^46^, Golgi staining and electron microscopy revealed decreased dendritic spine density and reduced postsynaptic density (PSD) thickness in hippocampal CA1 pyramidal neurons of LUZP1-cKO mice compared with control littermates (Figures S2C-S2F). Furthermore, loss of LUZP1 significantly increased primary cilium length relative to control littermates in hippocampal CA1 pyramidal neurons (Figures 2H and 2I). Although gross hippocampal development appeared preserved, membrane-associated acetylated α-tubulin levels were decreased in the LUZP1-cKO hippocampus relative to controls (Figure 2J and 2K). Together, these findings extend the cellular phenotypes to the brain hippocampus and support a requirement for LUZP1 in hippocampal structural maturation, ciliary homeostasis, and behavior.

**Figure 2.**
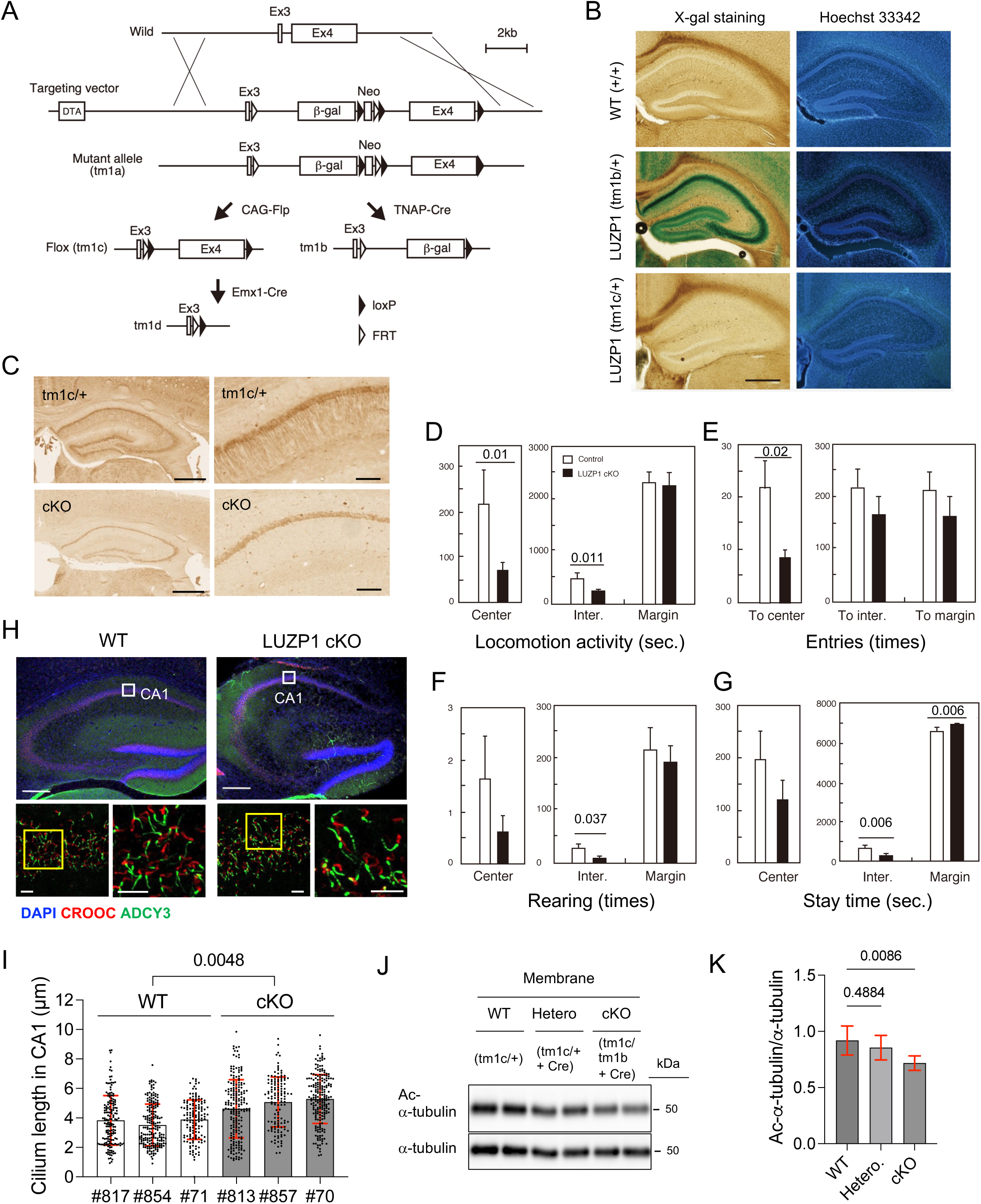
LUZP1 deficiency impairs locomotor behavior in mice. (A) Strategy for conditional *Luzp1* knockout (LUZP1-cKO) mouse generation. See Materials and Methods. (B and C) LacZ staining (B) and LUZP1 immunohistochemistry (C) in coronal brain sections. LUZP1 expression was detected by custom-made anti-LUZP1 immunoblotting. (D-G) Open-field test measuring locomotor activity (D), entries (E), rearing events (F), and time spent (G) in each compartment within a square arena. Control, n = 18; LUZP1-cKO, n = 15. (H) CA1 cilia in WT and LUZP1-cKO hippocampus. Upper: Hippocampal coronal 20 µm-thick section maximum intensity projection (MIP). Lower: Magnified images. 15 µm MIP. (I) Cilium length in CA1 hippocampus. n = 3, 2-month-old mice. (J and K) Membrane-associated acetylated α-tubulin in WT, heterozygous, and LUZP1-cKO hippocampus. Scale bars, 500 μm (B and C, left), 100 μm (C, right), 200 μm (H, upper), and 10 μm (H, lower). Data in K were pooled from three technical replicates from two mice per group. Data are presented as mean ± SD. Two-tailed unpaired t-tests (D-G and I). One-way ANOVA plus Dunnett’s multiple comparisons test (K). Exact P values are shown.

### LUZP1 interacts with HDAC6 and ATAT1 but preferentially couples to ATAT1 to promote α-tubulin acetylation

To define how LUZP1 regulates microtubule acetylation, we generated a *Luzp1* knockout HEK293T (LUZP1-KO HEK293T) cell line using CRISPR-Cas9 genome editing. Complete loss of LUZP1 protein was confirmed by immunoblot analysis (Figure 3A). Consistent with the phenotype observed in LUZP1-cKO mice, LUZP1 deficiency significantly reduced acetylated α-tubulin levels without altering total α-tubulin levels (Figure 3B). Re-expression of mouse LUZP1 in LUZP1-KO HEK293T cells increased α-tubulin acetylation (Figures 3C and 3D). Reduced acetylated α-tubulin levels were also detected in primary hippocampal and cortical neurons infected with AAV9-LUZP1 shRNA viruses (Figures 3E and S2G), indicating that LUZP1 positively regulates microtubule acetylation across multiple cellular contexts.

**Figure 3.**
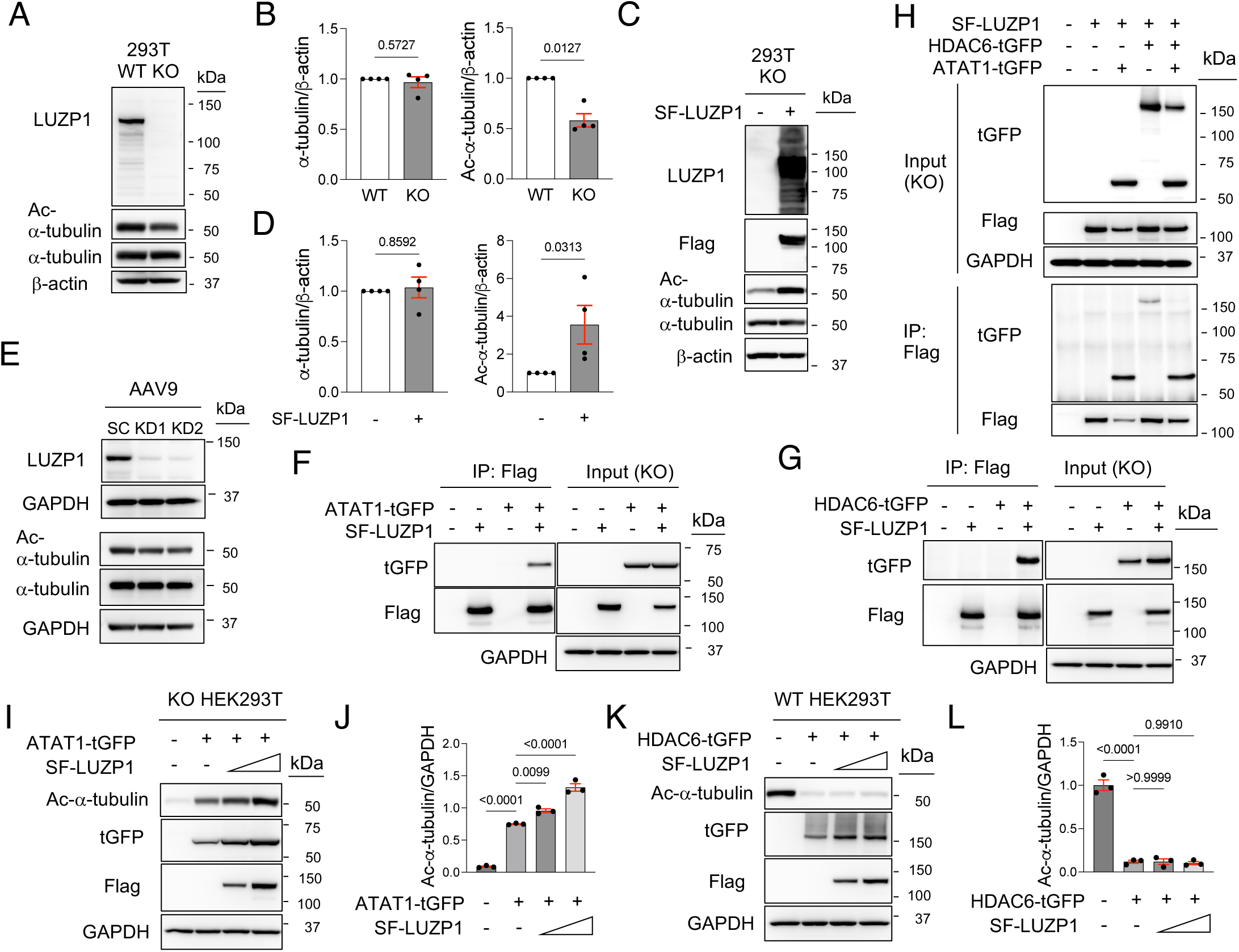
LUZP1 interacts with the microtubule-modifying enzymes HDAC6 and ATAT1 but selectively modulates ATAT1 activity toward α-tubulin. (A and B) Validation of CRISPR-Cas9-mediated LUZP1 knockout in HEK293T cells and quantification of acetylated α-tubulin. n = 4 independent experiments. (C and D) Rescue of α-tubulin acetylation by mouse LUZP1 re-expression in LUZP1-KO HEK293T cells. n = 4. (E) LUZP1 immunoblot in hippocampal neurons infected with AAV9-U6-shRNA. Hippocampal neurons were infected with AAV9-U6-shRNA virus at DIV9 and harvested at DIV17. (F–H) Co-immunoprecipitation assays testing LUZP1 interaction with ATAT1, HDAC6, or both. (I–L) Effects of LUZP1 on α-tubulin acetylation in cells coexpressing ATAT1 in LUZP1-KO HEK293T cells (I and J) or HDAC6 in WT HEK293T cells (K and L). n = 3. Data are presented as mean ± SEM. Ratio paired t-tests (B and D). Ordinary one-way ANOVA plus Tukey’s multiple comparisons test (J and L). Exact P values are shown.

We next asked whether LUZP1 interacts with ATAT1 and HDAC6, two core regulators of microtubule acetylation. Immunoprecipitation assays showed that LUZP1 interacted with both enzymes (Figures 3F and 3G) through its C-terminus, amino acid (aa) 636-1068 (Figure S3). Co-expression of LUZP1, ATAT1, and HDAC6 further showed that LUZP1 preferentially bound ATAT1 over HDAC6 (Figure 3H). Notably, LUZP1 enhanced ATAT1-mediated α-tubulin acetylation but did not affect HDAC6-mediated α-tubulin deacetylation (Figures 3I–3L). These data identify a selective LUZP1–ATAT1 module that promotes microtubule acetylation while leaving HDAC6 activity toward α-tubulin largely unchanged.

### ATAT1 and HDAC6 define separable LUZP1-dependent pathways for neurite and spine-cilia phenotypes

To test whether increasing microtubule acetylation is sufficient to rescue neuronal defects caused by LUZP1 deficiency, we overexpressed ATAT1 in LUZP1-knockdown hippocampal neurons. ATAT1 overexpression significantly restored axonal and dendritic length at DIV5 and DIV10, respectively (Figures 4A, 4B, and Figure S4). However, ATAT1 overexpression failed to rescue ciliary elongation or dendritic spine density at DIV12 and DIV20, respectively (Figures 4C, 4D, and Figure S4). We then treated LUZP1-knockdown hippocampal neurons with tubastatin A (TBA), a selective HDAC6 inhibitor that increases α-tubulin acetylation^47^. In contrast to ATAT1 overexpression, HDAC6 inhibition did not rescue axonal or dendritic length but significantly restored ciliary length and dendritic spine density in LUZP1-knockdown hippocampal neurons (Figures 4E–4H and Figure S5). These findings separate the LUZP1 phenotypes into distinct effector pathways: an ATAT1-dependent module that supports neurite extension and an HDAC6-sensitive pathway that governs ciliary growth and spine maturation.

**Figure 4.**
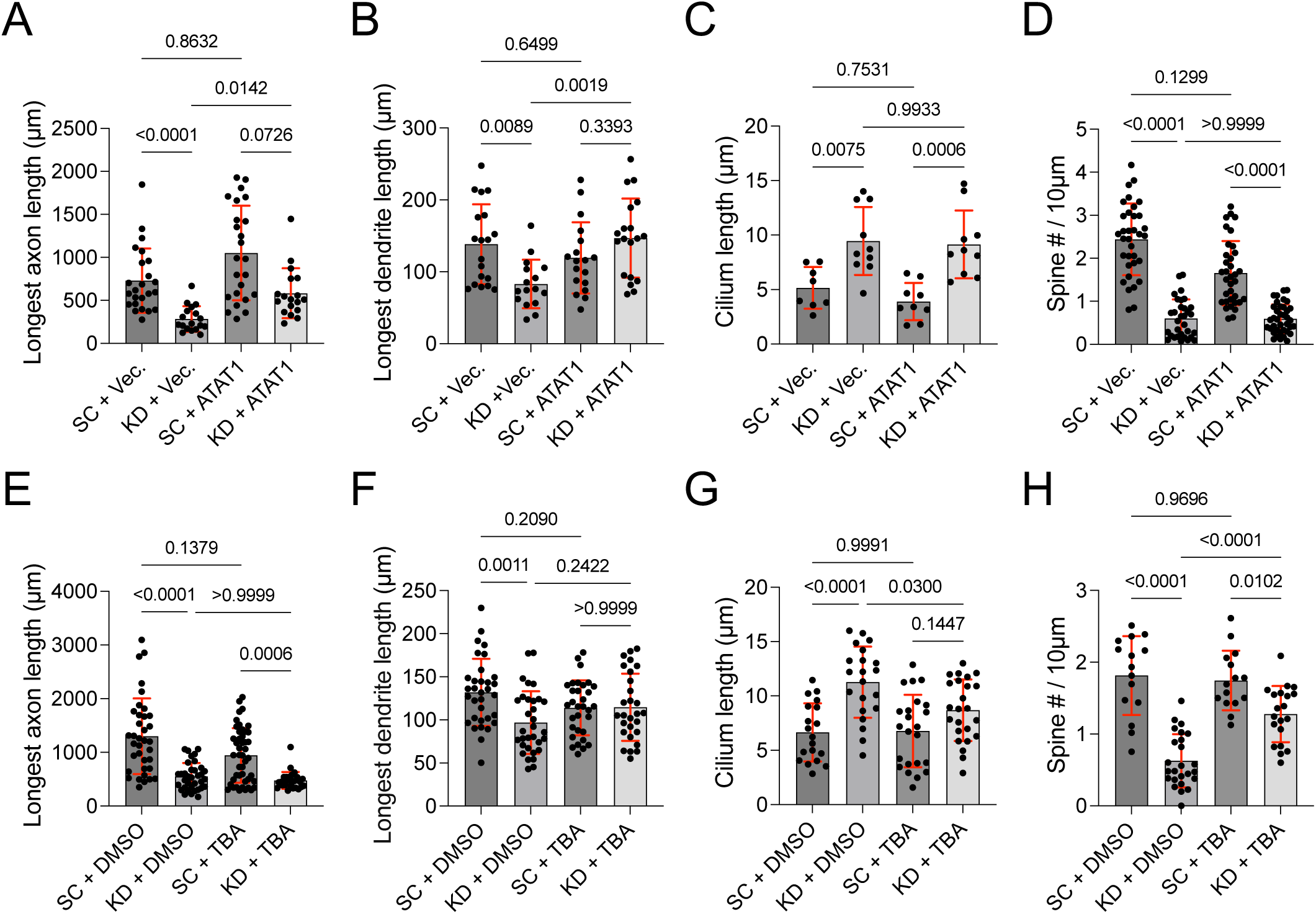
Differential roles of ATAT1 and HDAC6 in regulating neurite and ciliary growth and dendritic spine maturation in LUZP1-depleted hippocampal neurons. (A–D) ATAT1 rescue experiments in LUZP1-knockdown hippocampal neurons assessing axon length (A), dendrite length (B), cilium length (C), and spine density (D). (E–H) Parallel HDAC6 inhibition experiments assessing the same readouts. Neurons were transfected at DIV3, DIV7, DIV9, or DIV17 and fixed at DIV5, DIV10, DIV12, or DIV20, respectively. Each dot represents one neuron (A–C and E–G) or one dendritic branch segment from an individual neuron (4–6 neurons per condition; D and H). Data in E and F are from one representative experiment; two independent experiments gave similar results. Data in A–D, G, and H were pooled from two independent experiments. Data are presented as mean ± SD. Kruskal-Wallis test with Dunn’s multiple-comparisons test in A and E; one-way ANOVA with Dunnett’s multiple-comparisons test in B–D and F–H. Exact P values are shown.

### LUZP1 restrains HDAC6-dependent CTTN deacetylation to support spine maturation and limit ciliary growth

Because LUZP1 did not inhibit HDAC6-dependent deacetylation of α-tubulin, we asked whether HDAC6 acts through an alternative substrate to control dendritic spine maturation and ciliary growth. We therefore focused on CTTN, a known HDAC6 substrate^12^, in LUZP1-knockdown hippocampal neurons. Co-immunoprecipitation assays showed that LUZP1 interacts with CTTN (Figure 5A) and that CTTN preferentially associated with an N-terminal LUZP1 fragment (aa 1–437) (Figure S6). LUZP1 deficiency reduced CTTN acetylation, whereas LUZP1 re-expression markedly increased it (Figure 5B). Notably, acetylated CTTN was barely detectable in pull-down fractions (Figure 5A). CTTN also interacted with both LUZP1 and HDAC6 (Figure 5C), supporting a role for LUZP1 in restraining HDAC6-dependent CTTN deacetylation. Consistent with this model, LUZP1 suppressed HDAC6-mediated deacetylation of CTTN (Figure 5D and 5E), suggesting that excessive CTTN deacetylation contributes to the LUZP1-deficient phenotype. To test this directly, we overexpressed acetylation-mimetic CTTN-7KQ and deacetylation-mimetic CTTN 7KR mutants^48^. CTTN-7KQ, but not wild-type (WT) CTTN or CTTN-7KR, restored ciliary length and spine density in LUZP1-knockdown hippocampal neurons (Figures 6A–6D). By contrast, CTTN-7KR increased filopodia formation (Figures 6C and 6D). Consistently, CTTN-7KQ, but not WT CTTN or CTTN-7KR, restored PSD95 localization to dendritic shafts and spines (Figure 6C). Together, these findings identify CTTN acetylation as a key downstream effector through which LUZP1 limits ciliary growth and promotes spine maturation.

**Figure 5.**
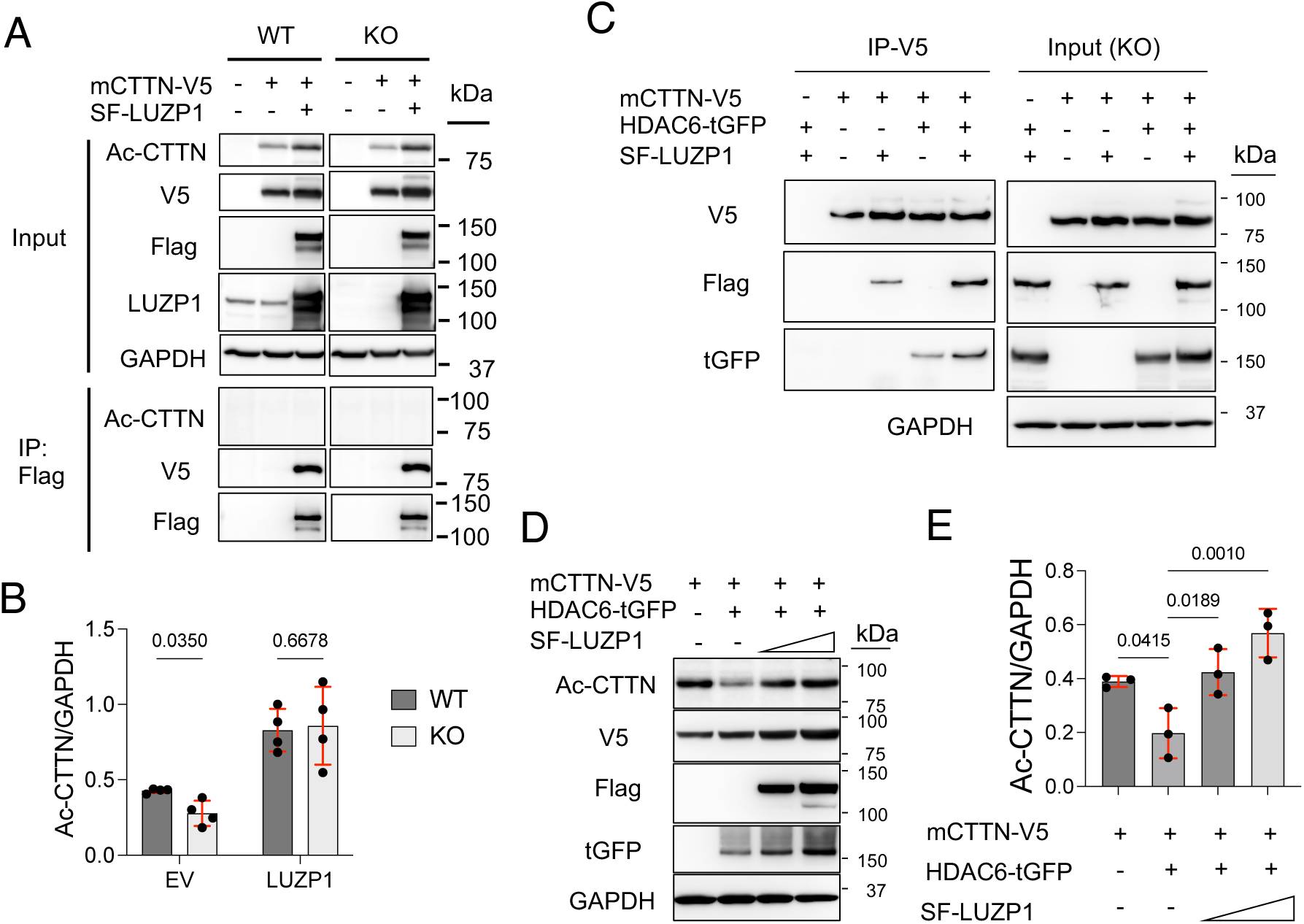
LUZP1 suppresses HDAC6-mediated CTTN acetylation. (A and B) CTTN co-immunoprecipitation and quantification of acetylated CTTN (AcCTTN) in WT or LUZP1-KO HEK293T cells. n = 4 independent experiments. (C–E) HDAC6–CTTN–LUZP1 interaction and effects of LUZP1 on CTTN acetylation. n = 3 independent experiments. Data are presented as mean ± SD. Two-way ANOVA with Šídák’s multiple-comparisons test in B; one-way ANOVA with Dunnett’s multiple-comparisons test in E. Exact p values are shown.

**Figure 6.**
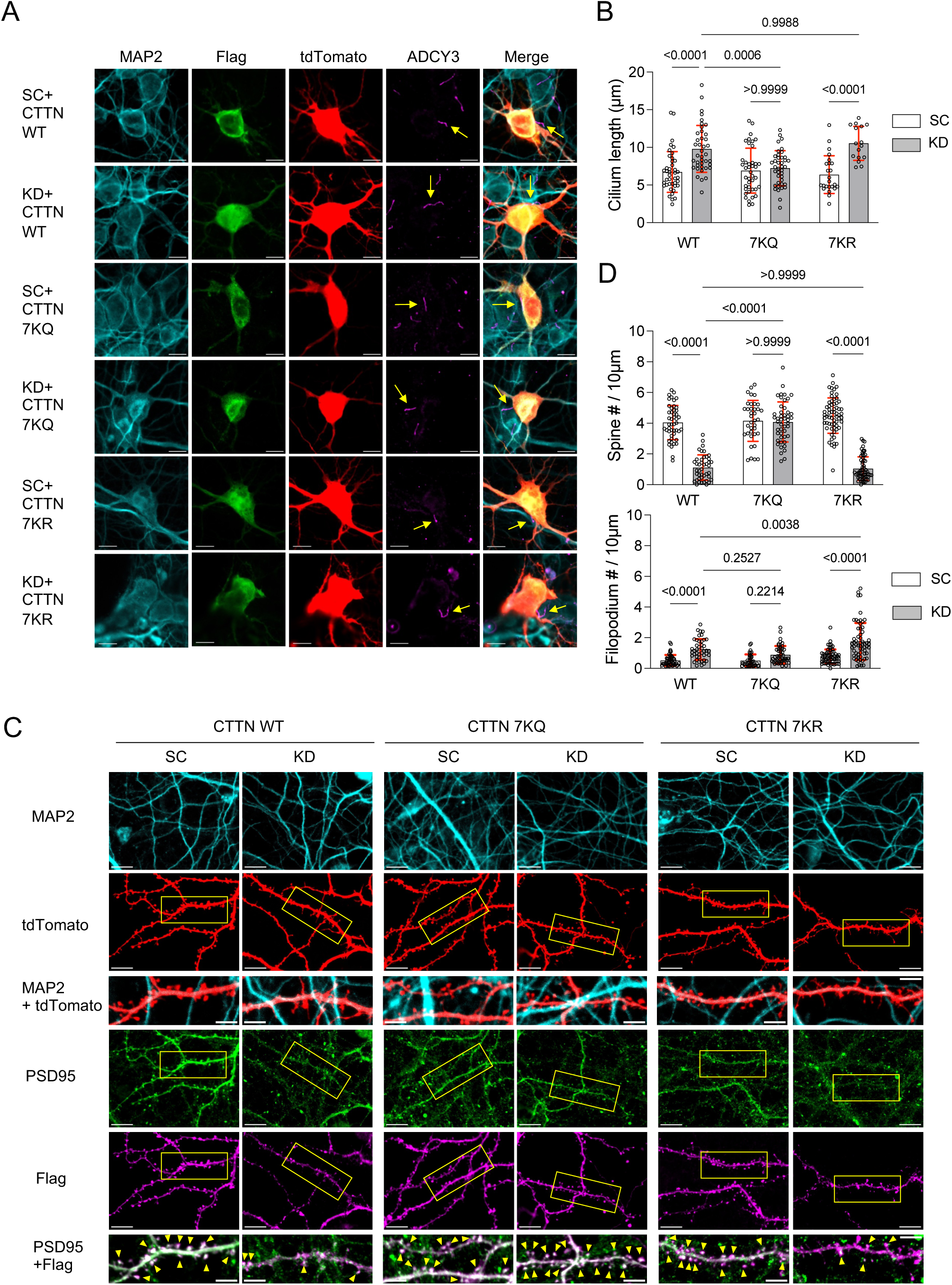
Acetyl-mimicking CTTN restores the abnormal ciliary length and spine maturation. (A and B) Cilia images and length quantification in hippocampal neurons expressing KD1 together with CTTN-WT, CTTN-7KR, or CTTN-7KQ. (C and D) Spine images and quantification after expression of human CTTN WT or the indicated mutants. Yellow arrowheads indicate colocalization of CTTN and PSD95 in tdTomato^+^ spines. Each dot represents one neuron (B) or a dendritic branch segment from an individual neuron (5–7 neurons per condition; D). Data in B and D are from one representative experiment; two independent experiments gave similar results. Scale bars: 10 µm (A and C). Two-way ANOVA plus Šídák’s multiple comparisons test (B and D). Data are presented as mean ± SD. Exact P values are shown.

### LUZP1 limits autophagy-dependent OFD1 degradation to constrain ciliary elongation

Autophagy is a known regulator of ciliogenesis^28,49^ and HDAC6-dependent CTTN deacetylation promotes autophagosome–lysosome fusion^50,51^. We therefore hypothesized that LUZP1 regulates ciliogenesis by controlling HDAC6-mediated CTTN acetylation, which in turn modulates autophagosome–lysosome fusion (Figure 7A). To test this hypothesis, we first examined autophagic flux in LUZP1-KO HEK293T cells by measuring LC3-II, an autophagosome marker, expression levels. Immunoblot analysis revealed increased autophagic flux in LUZP1-KO HEK293T cells (Figures 7B and 7C). In primary hippocampal neurons, LUZP1 knockdown similarly reduced the steady-state number of LC3-positive autophagosomes (Figures 7D and 7E). However, pharmacological suppression of autophagy caused a significantly greater accumulation of LC3-positive autophagosomes in LUZP1-knockdown neurons than in control neurons (Figure 7F). LUZP1 knockdown also reduced p62 levels, further supporting enhanced autophagic flux (Figure 7L, 7N and S7E). Notably, autophagy suppression reversed the ciliary elongation phenotype observed in LUZP1-knockdown hippocampal neurons and LUZP1-KO HEK293T cells (Figures 7G, 7H and S7A-C), indicating that enhanced autophagy directly drives aberrant ciliogenesis in the absence of LUZP1. By contrast, autophagy inhibition failed to rescue dendritic spine defects in LUZP1-knockdown hippocampal neurons (Figure 7I), suggesting that LUZP1-dependent spine maturation is mechanistically separable from its effects on ciliary homeostasis. Together, these results place autophagy downstream of the LUZP1–HDAC6–CTTN pathway in the control of ciliary growth.

**Figure 7.**
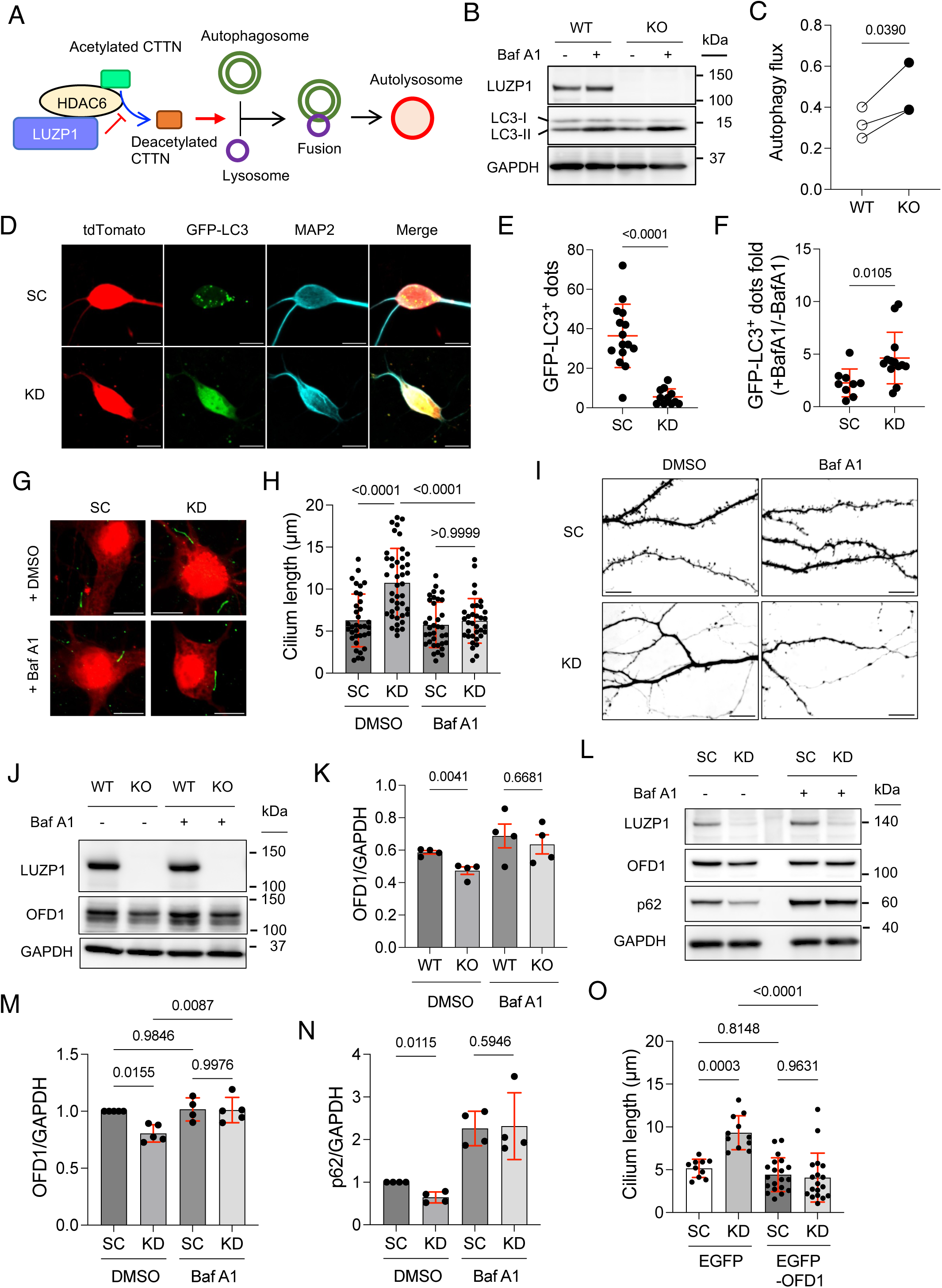
Autophagic OFD-1 degradation promotes ciliary elongation under LUZP1-deficient conditions. (A) Model for LUZP1-dependent regulation of autophagy through HDAC6–CTTN signaling. (B and C) LC3-II immunoblot and autophagic flux analysis in WT and LUZP1-KO HEK293T cells. Baf A1: Bafilomycin A1. “–” in the Baf A1 panel indicates DMSO-treated cells. LC3-II protein levels were quantified densitometrically and normalized to the loading control. Autophagic flux was calculated by subtracting normalized LC3-II values in DMSO-treated samples from those in Baf A1-treated samples. n = 3 independent experiments. (D) LC3-GFP^+^ puncta images in primary hippocampal neurons. SC and KD1 shRNA vectors were cotransfected with GFP-LC3-expressing vector at DIV7 and fixed at DIV12. (E) Quantification of GFP-LC3^+^ puncta from (D). (F) GFP-LC3 puncta analysis in control and LUZP1-knockdown hippocampal neurons with or without Baf A1. KD1 shRNA vectors were cotransfected with GFP-LC3-expressing vector at DIV8. Neurons were treated with Baf A1 for 6 hours prior to fixation at DIV13. Fold increase represents GFP-LC3^+^ puncta in Baf A1-treated (+Baf A1) groups normalized to the mean GFP-LC3^+^ puncta in DMSO-treated (-BafA1) groups. (G and H) Cilia analysis in Baf A1-treated LUZP1-knockdown hippocampal neurons. KD1 shRNA vectors were transfected into hippocampal neurons at DIV9, followed by Baf A1 treatment for 6 hours at DIV13 before fixation. Cilium length was quantified. (I) Spine images in Baf A1-treated LUZP1-knockdown hippocampal neurons. KD1 shRNA vectors were transfected into hippocampal neurons at DIV17, followed by Baf A1 treatment for 6 hours at DIV20 before fixation. (J and K) OFD1 immunoblot and quantification in WT and LUZP1-KO HeLa cells. n = 4. (L–N) OFD1 and p62 immunoblots and quantification in control and LUZP1-knockdown hippocampal neurons. Hippocampal neurons were infected with AAV9-U6-shRNA virus carrying scramble or KD1 sequences at DIV9 and harvested at DIV18. Neurons were also treated with Baf A1 for 6 hours before harvest. (O) Cilium length in control and LUZP1-knockdown neurons expressing EGFP-OFD1. Scale bars, 10 μm (D, G, and I). Each dot represents one neuron (E, F, H, and O). Data in E, and I are from one representative experiment; two independent experiments gave similar results. Data in F and O were pooled from two independent experiments, data in H from three, data in M from three KD1 and two KD2, and data in N from two KD1 and two KD2 experiments. Data are presented as means ± SD. Ratio paired t-tests (C). Two-tailed unpaired t-tests (E). Mann-Whitney test (F). Kruskal-Wallis plus Dunn’s multiple comparisons test (H). One-way ANOVA plus Tukey’s multiple comparisons test (O). Paired t test (K, M and N). Exact P values are shown.

Autophagy promotes primary ciliogenesis by degrading oral-facial-digital syndrome 1 (OFD1)^28,49^, a ciliogenesis suppressor. To further define the link among LUZP1, autophagy, and ciliogenesis, we generated a *Luzp1* knockout HeLa (LUZP1-KO HeLa) cell line, which provides robust OFD1 expression^52^, using CRISPR-Cas9 genome editing (Figure S7D). LUZP1-KO Hela cells reproduced the reduction in acetylated α-tubulin observed in LUZP1-KO HEK293T cells (Figure S7D). OFD1 degradation was accelerated in LUZP1-KO HeLa cells relative to controls (Figures 7J and 7K). Consistently, OFD1 levels were lower in LUZP1-knockdown hippocampal neurons, and autophagy inhibition with bafilomycin A1 restored OFD1 levels (Figures 7J–7M and S7E). Finally, OFD1 overexpression rescued the aberrant ciliogenesis phenotype in LUZP1-knockdown hippocampal neurons (Figure 7O). These findings identify autophagy-dependent OFD1 turnover as a downstream mechanism through which LUZP1 constrains ciliary elongation.

### Activated CaMKIIα selectively strengthens LUZP1 interactions with HDAC6 and CTTN

Because Ca^2+^ influx through postsynaptic NMDARs activates CaMKII to regulate F-actin organization in dendritic spines^40^, we investigated whether CaMKII phosphorylates LUZP1 and thereby alters its interactions with CTTN, HDAC6, and ATAT1. LUZP1 exhibited basal serine/threonine phosphorylation; however, co-expression of full-length CaMKIIα did not further increase this signal (Figure 8A). By contrast, a constitutively active truncated CaMKIIα (tCaMKIIα) enhanced LUZP1 phosphorylation relative to the kinase-dead K42R mutant (Figure 8B), suggesting that CaMKII activation promotes LUZP1 phosphorylation. Unexpectedly, both full-length CaMKIIα and its truncated variants interacted with LUZP1 (Figures 8A and 8B). Immunostaining revealed that CaMKIIα colocalized with LUZP1 along MAP2-positive dendritic microtubules and was enriched within dendritic spines (Figures S8A and S8B). Following LUZP1 knockdown, the distribution of CaMKIIα within MAP2-positive dendrites became more diffuse and exhibited reduced colocalization with dendritic microtubules, possibly reflecting alterations in the dendritic cytoskeletal organization. In addition, the presence of CaMKIIα within dendritic spines was reduced (Figures S8A and S8B). Notably, activated tCaMKIIα selectively enhanced LUZP1 interactions with CTTN and HDAC6, whereas the LUZP1–ATAT1 interaction remained unchanged (Figures 8C–8F). These findings identify LUZP1 as an activity-responsive effector of CaMKII signaling and place CaMKIIα upstream of the HDAC6–CTTN branch of the LUZP1 pathway.

**Figure 8.**
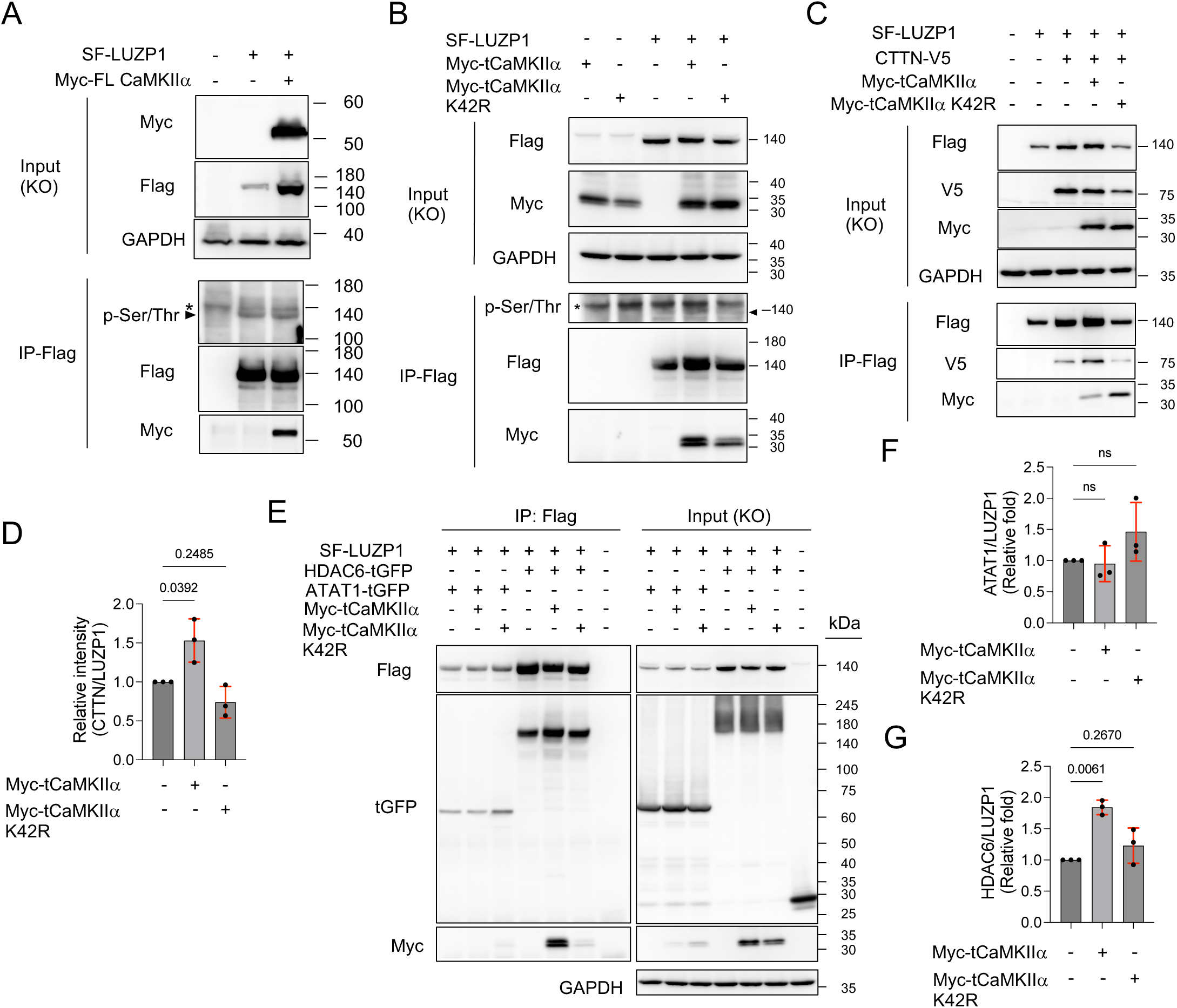
Activated CaMKIIα selectively enhances LUZP1 interactions with HDAC6 and CTTN. (A) Co-immunoprecipitation assay testing interaction of LUZP1 with full-length CaMKIIα. Arrowhead indicates phospho-LUZP1 and asterisk indicates non-specific binding proteins. (B) Co-immunoprecipitation assay testing interaction of LUZP1 with truncated CaMKIIα variants. Arrowhead indicates phospho-LUZP1 and asterisk indicates non-specific binding proteins. (C and D) Effects of truncated CaMKIIα variants on the LUZP1–CTTN interaction. n = 3 independent experiments. (E-G) Effects of truncated CaMKIIα variants on LUZP1 interaction with ATAT1 or HDAC6. n = 3 independent experiments. RM one-way ANOVA plus Dunnett’s multiple comparisons test (D, F and G). Data are presented as mean ± SD. Exact P values are shown.

## Discussion

Neurons continuously remodel specialized subcellular compartments through coordinated control of cytoskeletal architecture and organelle homeostasis. Although acetylation-dependent regulation of microtubule dynamics, actin organization, and autophagy has been implicated in neuronal development and plasticity, how these processes are integrated across maturation states has remained unclear. Here, we identify LUZP1 as a signaling organizer that links cytoskeletal acetylation to autophagy in neurons. Our data support a model in which LUZP1 governs neurite extension, dendritic spine maturation, and primary ciliogenesis through mechanistically separable modules involving ATAT1-dependent microtubule acetylation and suppression of HDAC6-mediated CTTN deacetylation. We further showed that LUZP1 engages CaMKIIα and that activated CaMKIIα selectively strengthens LUZP1 interactions with CTTN and HDAC6. This activity-responsive complex is consistent with a model in which neuronal activity tunes CTTN acetylation homeostasis to stabilize F-actin architecture in dendritic spines and soma. In parallel, loss of LUZP1 accelerated autophagy-dependent OFD1 degradation, thereby linking autophagic regulation to ciliogenesis. Together, these findings position LUZP1 as a signaling hub that coordinates cytoskeletal remodeling with autophagy during neuronal plasticity (Figure 9).

**Figure 9.**
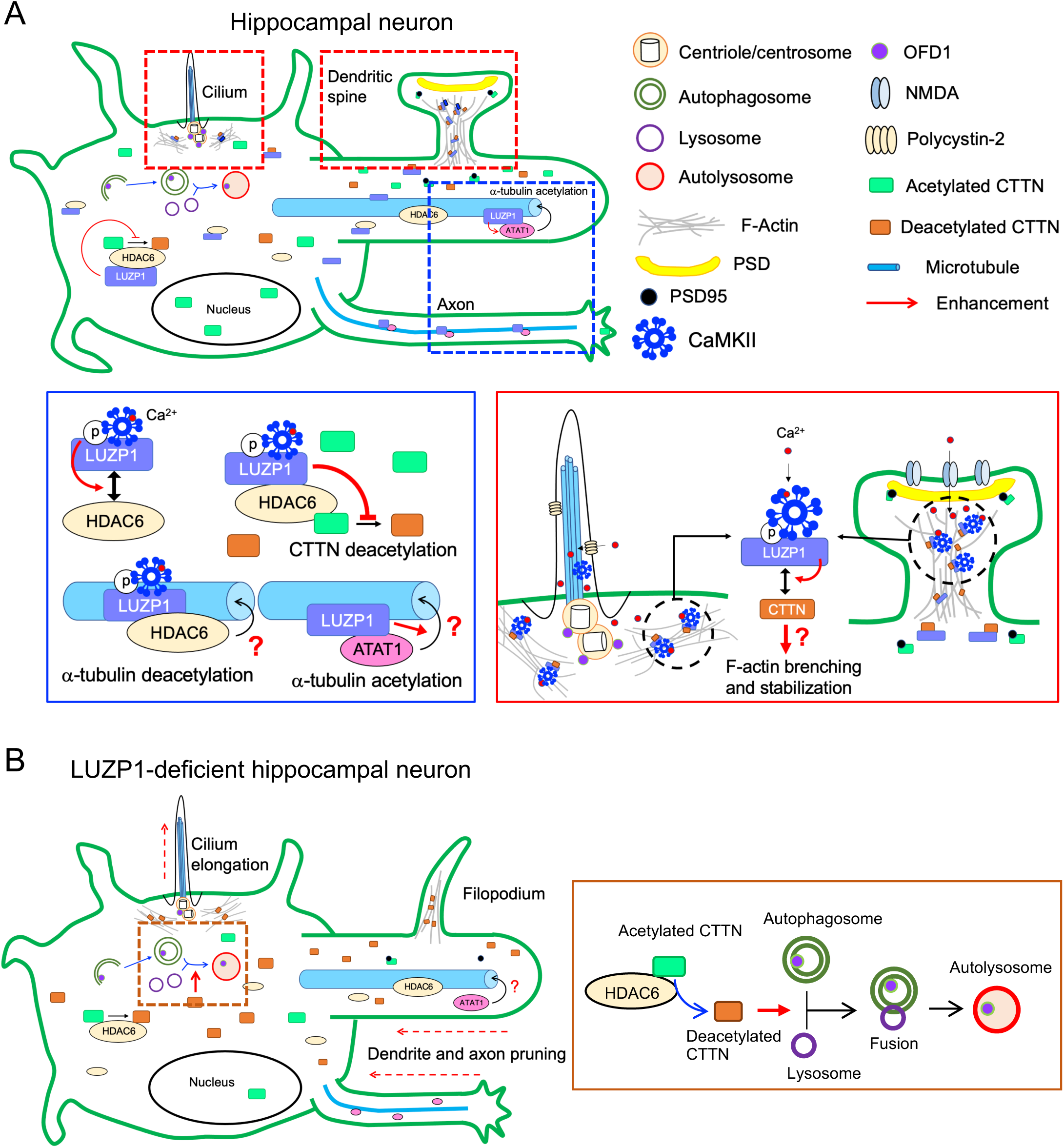
A schematic model of LUZP1 coordinating compartment-specific cytoskeletal remodeling in hippocampal neurons. (A) LUZP1 serves as a multifunctional regulator in hippocampal neurons, coordinating cytoskeletal dynamics, protein acetylation, and autophagy. LUZP1 promotes ATAT1-dependent α-tubulin acetylation, restrains HDAC6-dependent CTTN deacetylation, and limits autophagy-dependent OFD1 degradation. During neuronal activation (red and blue boxes), Ca^2+^ influx into dendritic spines and cilia through NMDA receptors and polycystin-2, respectively, activates CaMKII. CaMKIIα activation strengthens LUZP1 interaction with CTTN and HDAC6, but not ATAT1. Our findings propose a CaMKII–LUZP1 pathway that controls proper F-actin network organization and influences actin branching and cytoskeletal architecture. (B) Loss of LUZP1 disrupts this coordinated regulation, leading to decreased α-tubulin and CTTN acetylation and enhanced autophagy, ultimately affecting neurite formation, dendritic spine maturation, and primary cilia elongation in hippocampal neurons.

α-Tubulin acetylation is a hallmark of stable microtubules and a key determinant of neurite extension and neuronal polarization^5^. ATAT1 is the principal acetyltransferase responsible for this modification, yet the mechanisms governing its access to the microtubule lumen remain incompletely defined *in vivo*^53^. Our data identified LUZP1 as a microtubule-associated factor that engages ATAT1 and enhances its acetyltransferase activity toward α-tubulin. Given that LUZP1 directly associates with microtubules^37^, it may act as a scaffold that positions ATAT1 at sites permissive for luminal entry or enriches ATAT1 on stable microtubule subsets within extending neurites. In this framework, LUZP1 would function not simply as a binding partner, but as a spatial organizer of ATAT1-dependent microtubule acetylation during neuronal morphogenesis.

A central advance of this study is the identification of distinct acetylation-dependent LUZP1 modules across neuronal maturation. Although HDAC6 is a principal α-tubulin deacetylase, our data indicate that LUZP1 does not measurably restrain HDAC6-mediated tubulin deacetylation. Instead, pharmacological HDAC6 inhibition selectively normalized ciliary elongation and restored dendritic spine density in LUZP1-knockdown neurons without rescuing early neurite outgrowth defects. These results support a model in which LUZP1 constrains ciliary growth and promotes spine maturation primarily by limiting HDAC6-dependent deacetylation of CTTN, rather than by controlling α-tubulin acetylation. More broadly, these findings place LUZP1 upstream of separable acetylation-dependent effector pathways that are deployed across distinct neuronal compartments and developmental stages.

CTTN is a core organizer of dendritic spine actin architecture. Acetylation-mimetic CTTN mutants enhance PSD95 clustering through Shank-dependent interactions^11^, whereas deacetylation-mimetic mutants show increased F-actin binding affinity^12^. In this context, CaMKIIα is well positioned to couple synaptic activity to structural remodeling: activated CaMKIIα strengthens the LUZP1–HDAC6 interaction, and LUZP1 suppresses HDAC6-mediated CTTN deacetylation, suggesting that CTTN acetylation homeostasis functions as a checkpoint linking actin remodeling to coordinated changes in spine structure during LTP. Consistent with this model, LUZP1 preferentially associated with deacetylated CTTN, and CaMKIIα activation further enhanced this interaction, supporting the existence of an activity-responsive LUZP1–CTTN complex that stabilizes F-actin architecture during structural plasticity. Alternatively, by maintaining CTTN acetylation, it is likely that LUZP1 influences spine morphology through regulation of PSD95 clustering dynamics^54^. Together, these findings argue that LUZP1 regulates dendritic spine maturation primarily through actin-dependent mechanisms, providing a mechanistic explanation for the filopodial phenotype in LUZP1-depleted hippocampal neurons and for the inability of ATAT1 overexpression to rescue spine defects.

Our findings further suggest that LUZP1 operates within an activity-responsive actin regulatory module that influences ciliogenesis. This idea is supported by studies showing that Ca^2+^ signaling components, including polycystin-2 and CaMKII, localize to cilia^42,55^ and that modulation of Ca^2+^ levels alters ciliary length^55,56^. In this context, the CaMKIIα–LUZP1–CTTN axis may maintain an actin state that restricts ciliary growth. Loss of LUZP1 would disrupt this organization, relieving actin-mediated suppression and promoting ciliary elongation, consistent with our observations. At the same time, LUZP1 deficiency enhanced autophagy-dependent OFD1 degradation, providing a parallel mechanism that further promotes ciliogenesis. Together, these findings support a model in which activity-dependent CaMKII signaling converges on a LUZP1–CTTN–F-actin axis that interfaces with autophagy to control ciliary homeostasis. Although the precise contribution of CaMKII to neuronal ciliogenesis remains to be defined, this framework provides a mechanistic link between synaptic activity, actin dynamics, and primary cilium regulation.

*In vivo* analyses further support a role for LUZP1 in coordinating neuronal structural maturation and animal behavior. Neuron-specific deletion of *Luzp1* reduced dendritic spine density and PSD thickness in hippocampal CA1 pyramidal neurons, indicating that disruption of LUZP1-dependent structural regulation has consequences for synaptic architecture. Notably, LUZP1-cKO mice exhibited reduced levels of membrane-associated acetylated α-tubulin despite preserved gross hippocampal development, suggesting a selective defect in cytoskeletal acetylation rather than a generalized developmental abnormality. In addition to synaptic phenotypes, neurons in LUZP1-cKO mice displayed elongated primary cilia, extending our *in vitro* findings to the intact brain. Primary cilia are increasingly recognized as signaling compartments that influence synaptic development and plasticity through cilia–synapse communication^57^. In hippocampal pyramidal neurons, cilia are enriched for the ciliary serotonin receptor 6 (5-hydroxytryptamine receptor 6, 5-HTR6), and serotonergic axons directly contact these cilia, establishing 5-HTR6-dependent signaling that has been implicated in chromatin regulation^57^. Given the fact that 5-HTR6 is a pharmacological target of antipsychotics and antidepressants^58^, and altered 5-HTR6 expression has been reported in neuropsychiatric disorders^59^, ciliary elongation may perturb 5-HTR6-dependent ciliary signaling^33^. Such disruption could act in concert with impaired cytoskeletal acetylation to alter neuronal circuit output. Accordingly, the altered locomotor and exploratory behaviors observed in LUZP1-cKO mice may reflect combined disruption of synaptic maturation and cilia-dependent signaling rather than a single structural defect. Together, these findings support a model in which LUZP1-dependent coordination of cytoskeletal acetylation, spine maturation, and ciliary homeostasis contributes to neuronal circuit function by linking synaptic architecture with cilium-mediated signaling.

An initially counterintuitive finding is that LUZP1-knockdown neurons exhibit both reduced α-tubulin acetylation and elongated cilia. We propose that this apparent paradox reflects the dominant influence of autophagy-dependent mechanisms. Although reduced microtubule acetylation would be expected to impair ciliogenesis, enhanced autophagic degradation of OFD1 appears to override this effect, resulting in net ciliary elongation. This model underscores the importance of pathway integration in shaping developmental outcomes and highlights how perturbation of a single regulatory node can generate complex phenotypes.

In summary, our study identifies LUZP1 as a multifunctional signaling organizer that coordinates acetylation-dependent control of microtubules, actin dynamics, and autophagy to direct neuronal plasticity. By enhancing ATAT1 activity while restraining HDAC6-mediated CTTN deacetylation, LUZP1 enables compartment-specific cytoskeletal remodeling that supports neuronal development and plasticity. We further identify LUZP1 as an activity-responsive effector of CaMKII signaling. Together, these findings provide a mechanistic framework for how cytoskeletal acetylation and autophagy are integrated during neuronal maturation and how disruption of this coordination may broadly alter neuronal circuit formation and function.

### Limitations of the study

This study has several limitations. First, although our data place LUZP1 upstream of ATAT1-dependent α-tubulin acetylation, we do not yet know whether LUZP1 directly enhances ATAT1 catalysis through allosteric regulation or instead acts primarily by organizing ATAT1 within microtubule-associated compartments. Second, the mechanism by which LUZP1 selectively restrains HDAC6 activity toward CTTN while leaving α-tubulin deacetylation largely unaffected remains unresolved. Third, because LUZP1 is a microtubule-interacting protein, it may influence CaMKII translocation to microtubules during LTP, but this possibility was not tested directly. In addition, although we identify autophagy-dependent regulation of ciliogenesis downstream of LUZP1, we do not directly assess how altered ciliary structure affects ciliary signaling or neuronal output in LUZP1-deficient neurons. Future studies will be needed to determine whether pathways such as Hedgehog, Wnt, or GPCR signaling are altered under these conditions and how such changes shape neuronal physiology. Finally, many of our conclusions are based on *in vitro* neuronal systems. Further work *in vivo* will therefore be required to define how LUZP1-dependent control of microtubule and CTTN acetylation contributes to circuit organization and behavior.

## Materials and Methods

### Animals

Pregnant ICR mice were obtained from either the Animal Facility at the University of Osaka or SLC (Shizuoka, Japan). LUZP1 cKO mice were generated at the Animal Facility of the University of Fukui. Mice were housed under standard conditions on a regular light/dark cycle with ad libitum access to food and water. All animal procedures were approved by the Animal Care and Use Committees of the University of Osaka and the University of Fukui.

### Expression vectors and plasmid construction

The mouse *Luzp1* gene was cloned into the pCAG vector with either an N-terminal twin-Strep-Flag tag or a C-terminal 3×Flag tag. The pCMV6-hATAT1-tGFP expression vector was purchased from OriGene (RG208373). The *ATAT1* gene was further subcloned into the pCAG-Myc c1 vector using the BglII and XhoI restriction sites. The pcDNA3.1-mHDAC6-HA plasmid was kindly provided by Dr. Toshifumi Fukuda^60^. To generate the pCMV6-mHDAC6-tGFP expression vector, the HDAC6 DNA fragment was excised from pcDNA3.1-mHDAC6-HA with EcoRI and XhoI and inserted into the pCMV6-hATAT1-tGFP plasmid digested with BglII and XhoI to replace the *hATAT1* insert. The pcDNA3.1-HFF-human cortactin WT, 7KQ, and 7KR expression vectors were kindly provided by Dr. Minoru Yoshida^48^ (RIKEN, Wako, Japan). pCAG-mGFP-Actin was purchased from Addgene (plasmid #21948). The GFP-LC3 expression vector was kindly provided by Dr. Tamotsu Yoshimori. pCMV10-3xFLAG-OFD1 and pCDNA3.1-Myc-eGFP-OFD1 plasmids were kindly provided by Dr. Brunella Franco^61^. CaMKIIα WT and K42R plasmids were kindly provided by Dr. Min-Jue Xie, and the full-length and truncated variants (aa 1–270) were further cloned into the pCAG-Myc c1 vector^62,63^.

For mouse *Luzp1* knockdown experiments, two targeting sequences were used: KD1, 5’-TGAGAAACTTAAGACACAAAT-3’, and KD2, 5’-TGCGGTCTAGAGCAATTATAA-3’. Annealed oligonucleotides were inserted into the pLKO.1-puro vector (Addgene, plasmid #8453). The hPGK promoter-puromycin cassette was replaced with a CAG promoter-tdTomato cassette using the SacII and KpnI restriction sites to generate the pLKO.1-U6-shLUZP1-CAG-tdTomato vector. For the scramble control, the sequence 5’-CCTAAGGTTAAGTCGCCCTCG-3’ was used. The same sequences were also cloned into the pAAV-U6-zsGreen vector (AAVpro® Helper Free System, Takara) for AAV9-U6-shRNA virus production according to the manufacturer’s instructions. For knockout of the human *LUZP1* gene, the guide sequence 5’-GTAGCTTTATCTCCTCCAGT-3’ was inserted into the BbsI-digested px458 vector (Addgene, plasmid #48138).

### Cell culture and Transfection

LUZP1-knockout HEK293T and HeLa (Kyoto, RRID:CVCL_1922) cell lines were generated using CRISPR-Cas9 genome editing. Cells were transfected with px458-*LUZP1* guide vectors using Lipofectamine 3000 (Thermo Fisher Scientific). Twenty-four hours later, GFP^+^ cells were single-cell sorted using a BD FACSAria III cell sorter and cultured for at least 2 weeks to obtain sufficient cells for validation of LUZP1 deletion by immunoblot analysis. Cells were maintained in Dulbecco’s modified Eagle’s medium (DMEM) supplemented with 10% fetal bovine serum and penicillin/streptomycin (Thermo Fisher Scientific). For autophagy inhibition in cell lines, bafilomycin A1 (200 nM; Selleck) was applied 4–6 h before cell harvest. To examine primary cilia, equal numbers of WT or LUZP1-KO HEK293T cells were plated to confluence in regular culture medium for 24 h and then serum-starved in DMEM for an additional 24 h before fixation and staining.

Primary hippocampal neurons were isolated from E16.5 mouse brains. Hippocampi were digested with papain (Worthington Biochemical Corp.) and DNase I (Sigma-Aldrich), dissociated, and plated on polyethyleneimine-coated four-well chambered glass slides (Matsunami Glass Ind., Ltd.) or 12-well culture dishes (Corning Life Sciences). Hippocampal neurons were cultured in Neurobasal medium supplemented with B27 and 1× GlutaMAX (Thermo Fisher Scientific).

### Immunoblot, immunoprecipitation and immunofluorescence microscopy analysis

HEK293T cells were transfected with indicated expression vectors using Lipofectamine 3000 (Thermo Fisher Scientific Inc.) at a 1:1 ratio (DNA to Lipofectamine3000). Twenty-four hours later, immunoblot analysis was performed to assess protein expression levels. Immunoprecipitation assays were conducted using indicated antibodies with Pierce™ Protein G Magnetic Beads (Thermo Fisher Scientific Inc.) according to the manufacturer’s protocols.

For immunofluorescence assays, hippocampal neurons were transfected with indicated plasmids mixed with Lipofectamine2000 (Thermo Fisher Scientific Inc.) at a 1:1.5 to 1:2 ratio (DNA to Lipofectamine2000) for 45 minutes at specific developmental stages (DIV3, DIV7, DIV9, DIV15, and DIV17) depending on the experimental design. To inhibit HDAC6 activity, the selective HDAC6 inhibitor Tubastatin A (TBA, 200 nM, Selleck) was applied to neurons 24 hours after transfection. For autophagy inhibition in neurons, 100 nM bafilomycin A1 (Selleck), was applied to neurons 6 hours before fixation. Neurons or HEK293T cells were fixed with 4% paraformaldehyde in phosphate-buffered saline (PBS, 15 min, RT) and permeabilized with 0.1 % Triton X-100/PBS (15 min, RT) before staining for relevant markers overnight at 4°C. For Twin-Strep Tag labeling, Strep-Tactin^®^XT DY-488 and DY-649 (IBA Lifesciences) were used. The samples were mounted using Anti-Fade Fluorescence Mounting Medium – Aqueous, Fluoroshield (ab104135; Abcam) and imaged with an FV3000 confocal microscope (Olympus) operated using FV31S-SW. For imaging of CaMKIIα subcellular localization, the ZEISS Lattice SIM5 Super resolution microscope was used.

For immunofluorescence assays in mouse brain, brain sections were prepared from 8-weeks old WT and cKO mice. Mice were transcardially perfused with PBS, followed by a fixative containing 4% paraformaldehyde in PBS. Brains were removed and postfixed in the same fixative for 2 h at 4°C, followed by immersion in PBS overnight at 4°C. Coronal brain sections (50 μm thick) were cut using a vibrating microtome and processed as free-floating sections. Sections were permeabilized with 0.3 % Triton X-100/PBS (2 h, RT) before staining for relevant markers overnight at 4°C. The antibodies used in this study are listed in the supplementary table 1.

Axon length, dendrite length, dendritic spine density, and cilia length were analyzed using confocal microscopy and quantified with Fiji software^64^.

### Generation of *Luzp1* conditional knockout mice (cKO)

Two *Luzp1* ES cell lines (*Luzp1* F01 and A04, C57BL6N strain) were obtained from the UC Davis KOMP repository^65^. *Luzp1* chimeric mice were generated by ES cell microinjection into ICR fertilized eggs at Kyoto University. These chimeric mice were crossed with C57BL6N mice to establish *Luzp1* tm1a/+ mice. *Luzp1* tm1a/+ mice were crossed with *Tnap*-Cre mice to produce *Luzp1* tm1b/+ mice and with CAG-Flp mice to produce *Luzp1* tm1c/+ mice^65–67^. *Luzp1* tm1b/+ and tm1c/+ mice were backcrossed with C57BL6J mice for ten generations. *Luzp1* tm1b/+; *Emx*1-Cre mice were crossed with *Luzp1* tm1c/tm1c mice to generate *Luzp1* conditional knockout mice. Genotyping primer sequences are as follows: LUZP1-18099R: gcacacaatctccacccttt, LUZP1-10809F: tgaacctccctatccctcct, LUZP1-22081R: acgctcctagagccttctcc, LacZ-C-For: cggtcgctaccattaccagt, Cre-For: ccatctgccaccagccag, Cre-Rev: tcgccatcttccagcagg, Fabpi-500F: cctccggagagcagcgattaaaagtgtcag, Fabpi-R: tagagctttgccacatcacaggtcattcag.

### Golgi staining of hippocampal neurons

Brains were prepared from 8-week-old WT and cKO mice and were processed according to the FD Rapid GolgiStain Kit protocol (FD NeuroTechnologies, MD, USA). Brains were sectioned into 100 μm thick slices using a cryostat. Serial images were analyzed using ImageJ with the Extended Depth of Field plugin.

### Isolation of membrane fraction

Isolated hippocampal tissue was homogenized with 10 volumes of isotonic Sucrose-HEPES buffer (0.32 M sucrose, 10 mM HEPES pH 7.4, 2 mM EDTA, and protease inhibitor cocktail) using a Dounce homogenizer. The homogenate was centrifuged at 1,000 × g for 15 min, and the supernatant (S1 fraction) was transferred to a new tube. The S1 fraction was centrifuged at 20,000 × g for 45 min, and the supernatant (S2 fraction) was removed. The pellet (P2 fraction) was washed with Sucrose-HEPES buffer and dissolved in 0.1% SDS and 1% Triton X-100 in Sucrose-HEPES buffer.

### Generation of anti-LUZP1 antibody

The mouse LUZP1 peptide (amino acids 484–502) was synthesized and used to immunize rabbits to generate antiserum against LUZP1. To purify LUZP1 antibodies from the antiserum, GST-LUZP1 protein (amino acids 438–635) was expressed in *E. coli* and used to construct a purification column.

### Transmission electron microscope examination

Brain sections were prepared from 8-weeks old WT and cKO mice. Mice were transcardially perfused with PBS, followed by a fixative containing 2.5% glutaraldehyde and 2% paraformaldehyde in PBS. Brains were removed and postfixed in the same fixative for 2 h at 4°C, followed by immersion in 0.1 M phosphate buffer (PB) overnight at 4°C. Coronal brain sections (50 μm thick) were cut using a vibrating microtome and processed as free-floating sections. After sectioning, sections were washed several times in 0.1 M PB (pH 7.4) and postfixed with 1% osmium tetroxide in 0.1 M PB for 1 h on ice. Sections were then rinsed with distilled water and stained with 0.5% uranyl acetate in distilled water overnight at 4°C. Samples were dehydrated through a graded ethanol series (65%, 75%, 85%, 95%, and 100%) at room temperature, followed by dehydration with anhydrous ethanol and propylene oxide. Tissues were subsequently infiltrated with a 1:1 mixture of propylene oxide and epoxy resin overnight at room temperature and flat-embedded in Epon 812. After polymerization, ultrathin sections were cut using an Ultracut microtome (Reichert-Nissei, Leica Microsystems) and observed using a transmission electron microscope (H-7650, Hitachi, Tokyo, Japan). Electron micrographs were acquired at comparable magnifications under identical imaging conditions. PSDs were identified based on electron-dense structures apposed to presynaptic terminals. Quantitative analyses of PSD area, PSD length, and PSD thickness were performed using ImageJ software. All measurements were initially obtained using the original ImageJ output units and subsequently converted to physical units based on scale bar calibration. Calibration was performed using the scale bar embedded in each micrograph (500 nm), and all values were converted to nanometers (nm) or square nanometers (nm²) accordingly. Quantification was performed blind to genotype.

### Open-field test

Open-field experiments measured locomotor activity, number of rearing events and entries, and time spent in each compartment within a square arena (48 cm × 48 cm) (MELQUEST Co., Toyama, Japan). Mice were placed in the center of the arena and allowed free movement for 120 min while being tracked by the SCANET MV-40 automated tracking system (Noldus Information Technology, Wageningen, Netherlands).

### Statistical analysis

GraphPad Prism 9 (GraphPad software) was used for statistical analyses. The tests used for statistical analyses are described in the figure legends. p < 0.05 was considered to be significant.

## Acknowledgments

The authors thank Drs. James D. Sutherland, Brunella Franco, Minoru Yoshida, Toshifumi Fukuda, Takashi Nakakura, Koichi Ikuta, Min-Jue Xie, Ko Miyoshi, and Tokuichi Iguchi for providing experimental materials and technical help; Sota Watanabe and Thanapong Boonyaniwas for technical help; and Yasuhiko Sato for assistance with the ZEISS Lattice SIM5 super-resolution microscope. This work was supported by JSPS KAKENHI Grant Number JP26670089 (to K. K.), JP20500307 (to H. Y.), JP24K09667 (to Y. O.), JP23K05989 (to M. T.), and MEXT KAKENHI Grant Number JP25293043 and JP17H04014 (to M. S.).

## Author contributions

M. S. first conceived and supervised the project, then started the project with K. K. Later, C.-Y. T. joined, designed, and carried out the experiments, analyzed data and wrote the manuscript with K. K. and M. S. K. K. performed the animal experiments, prepared the materials, and helped to analyze data. M. T. performed the TEM experiments and helped to analyze data. Y. O. performed the ISH experiments and provided the materials. H. Y. performed some of the animal experiments.

## Data and materials availability

All other data needed to evaluate the conclusions in the paper are present in the paper or the Supplementary Materials.

## Declaration of interests

The authors declare no competing interests.

## Declaration of generative AI and AI-assisted technologies in the English editing

The authors used ChatGPT-5 and Copilot to assist with English editing and polishing. All content was reviewed and verified by the authors, who take full responsibility for the final manuscript.

## Supplementary Figure legends

**Figure S1.**
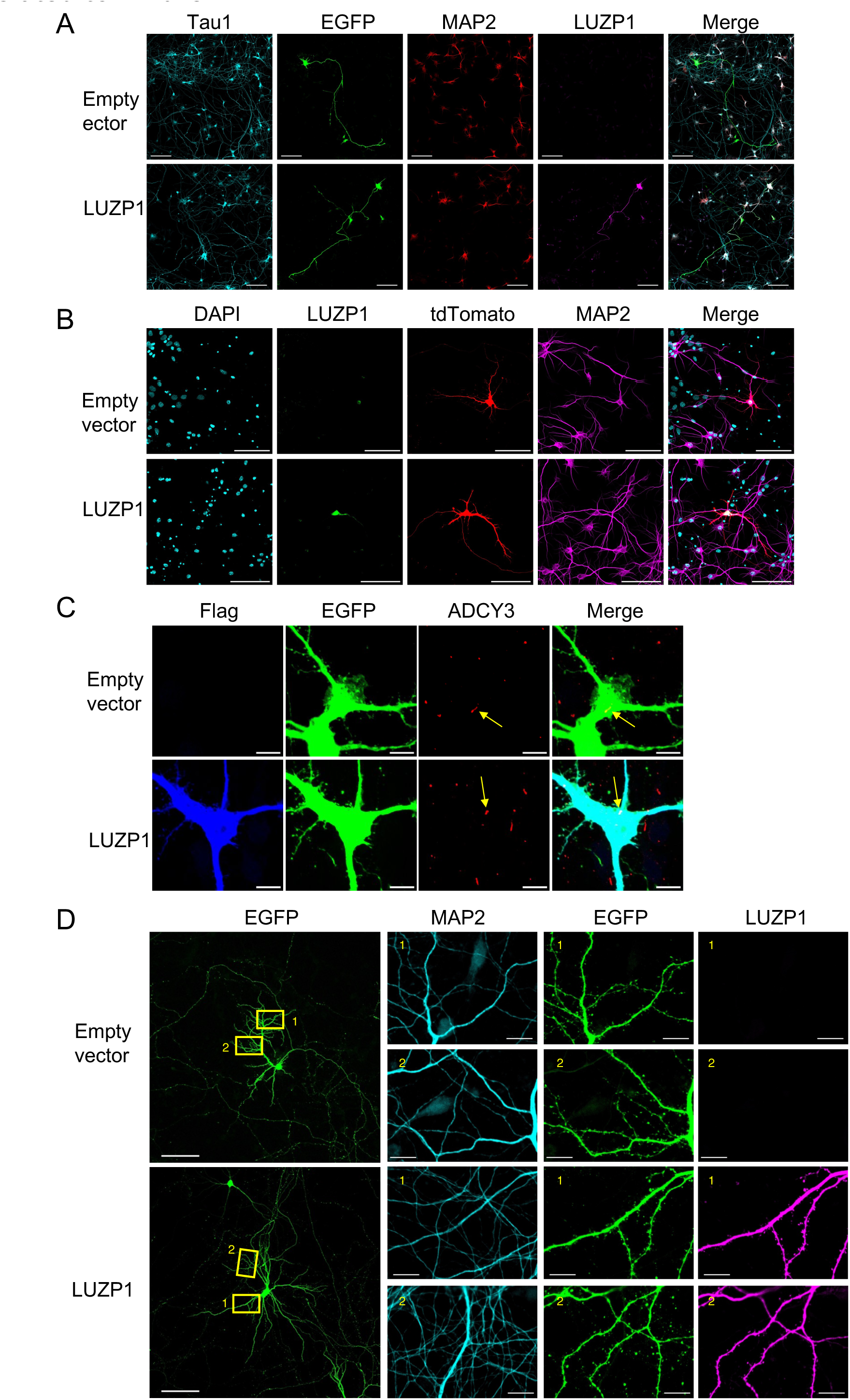
LUZP1 overexpression in hippocampal neurons does not alter axon, dendrite, or spine formation. Related to Figure 1. Representative images of axons, dendrites, and spines in hippocampal neurons overexpressing SF-LUZP1 at DIV3 (A), DIV7 (B), DIV9 (C) or DIV17 (D) and fixed at DIV5, DIV10, DIV12, or DIV20, respectively. Scale bars, 100 μm (A–D) and 10 μm (D, inset).

**Figure S2.**
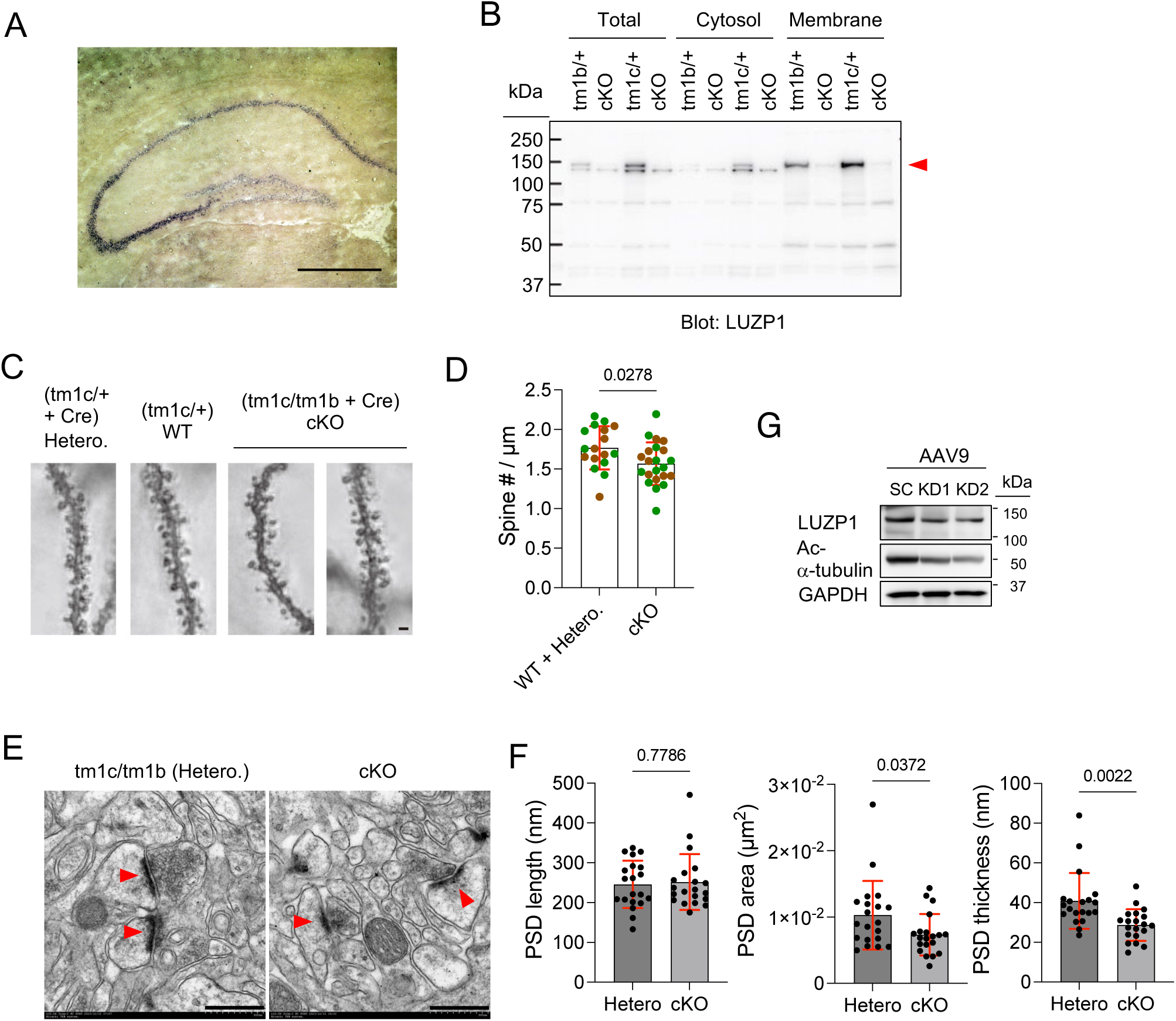
LUZP1 deficiency reduces dendritic spine density and postsynaptic density thickness. Related to Figure 2. (A) *In situ* hybridization for *Luzp1* in WT hippocampus. (B) LUZP1 immunoblot in total, cytosolic, and membrane fractions from WT, heterozygous (Hetero.), and LUZP1-cKO hippocampus. LUZP1 expression was detected by custom-made anti-LUZP1 immunoblotting. The red arrowhead indicates LUZP1. (C and D) Golgi staining of hippocampal neurons and spine density quantification *in vivo*. (E and F) Electron micrographs of dendritic spines and quantification of PSD length, area, and thickness. n = 1 mouse per group. (G) LUZP1 knockdown and acetylated α-tubulin expression in primary cortical neurons infected with AAV9-U6-shRNA at DIV16 and harvested at DIV26. The SC and KD sequences used in G were identical to those in Figure 1A. Scale bars, 500 μm (A), 1 μm (C), and 500 nm (E). Data are presented as means ± SD. Two-tailed unpaired t test in D and F. Exact p values are shown.

**Figure S3.**
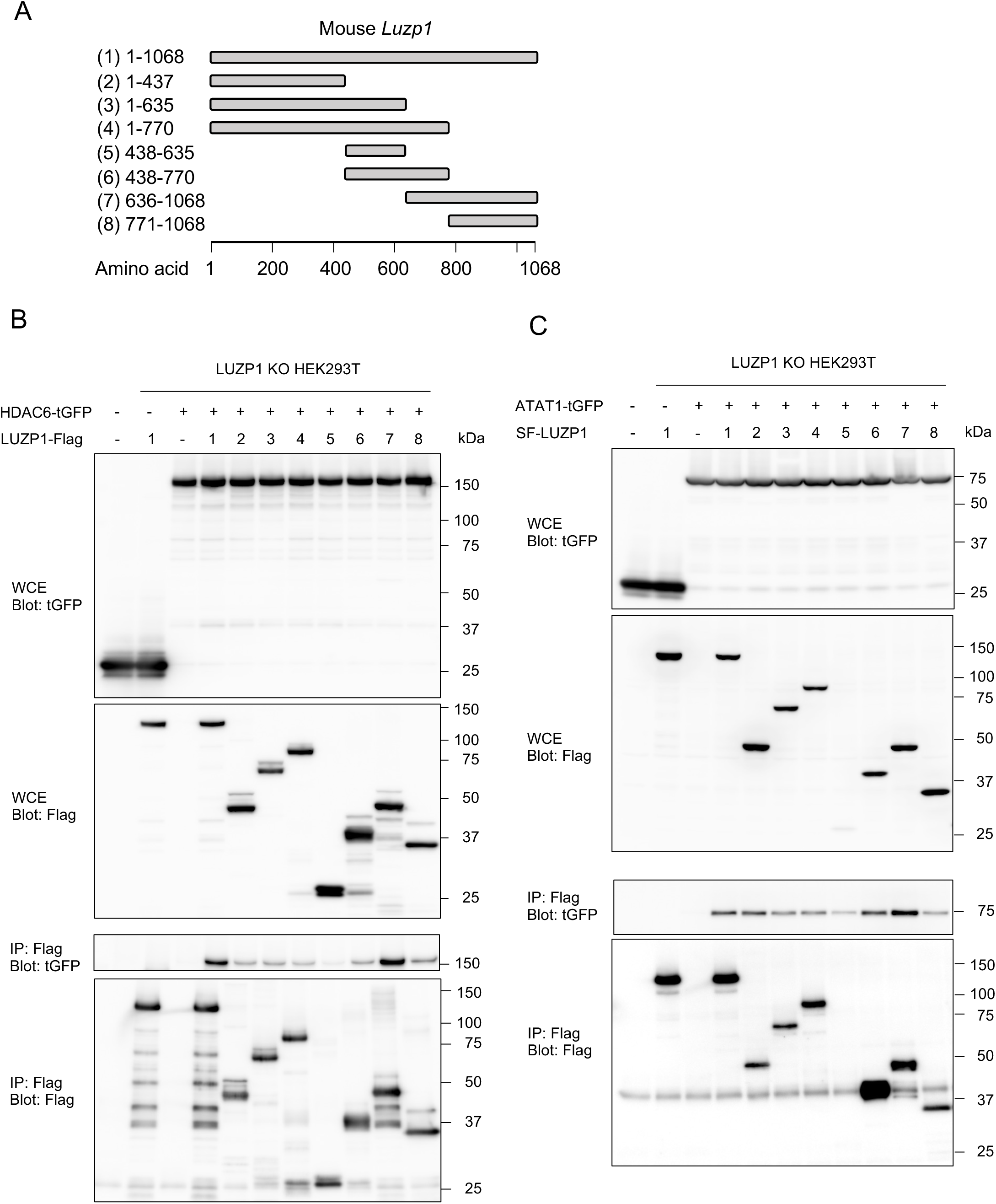
LUZP1 C-terminus interacts with both HDAC6 and ATAT1. Related to Figure 3. (A) Schematic of WT and truncated LUZP1 constructs. (B and C) Immunoprecipitation assays testing interaction of HDAC6 (B) or ATAT1 (C) with WT or truncated LUZP1 in LUZP1-KO HEK293T cells. Two independent experiments gave similar results.

**Figure S4.**
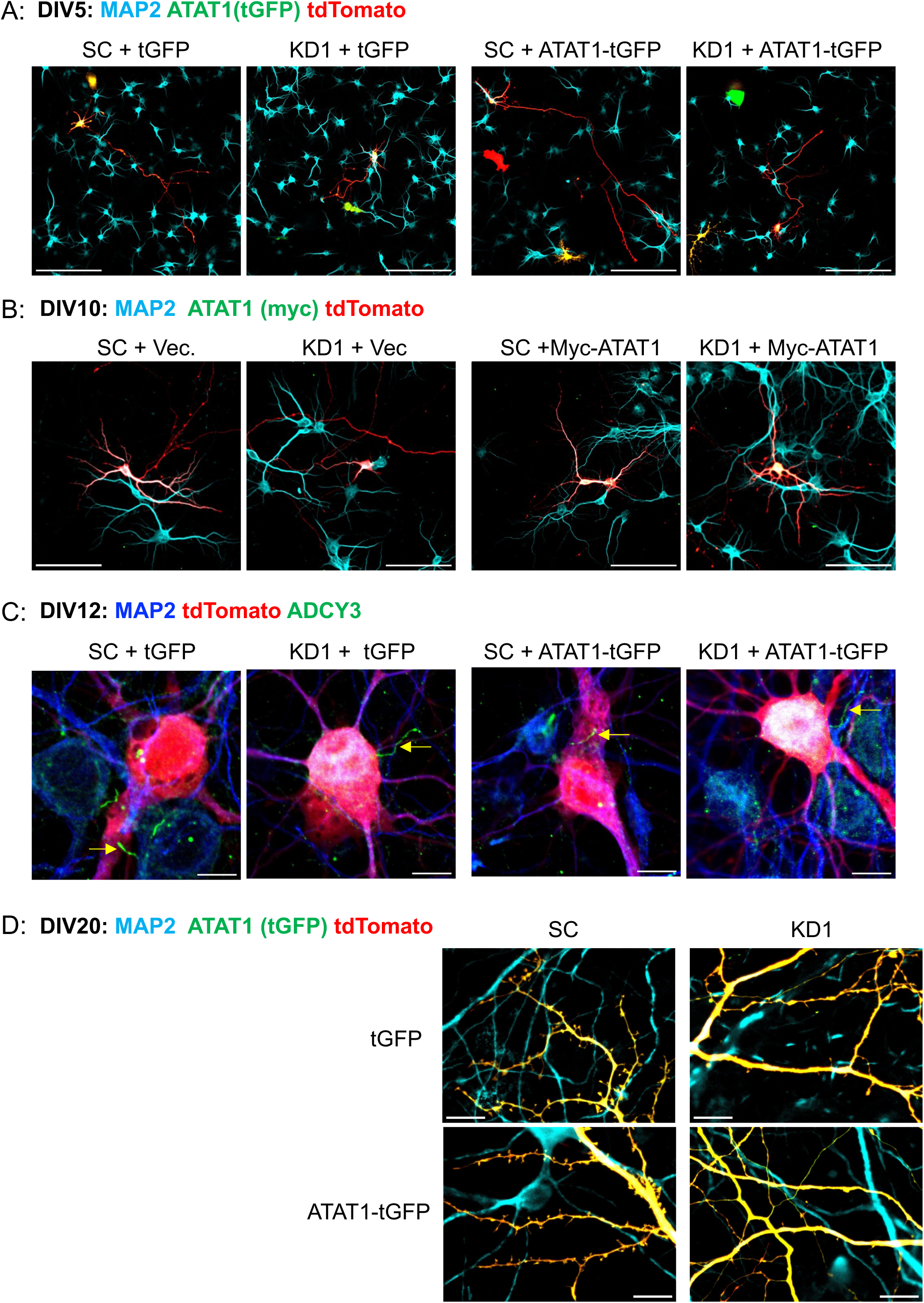
ATAT1 overexpression rescues axon and dendrite defects but not cilium and spine defects in LUZP1-depleted neurons. Related to Figure 4. Yellow arrowheads indicate ACDY3^+^ cilia. Scale bars, 100 μm (A and B) and 10 μm (C and D).

**Figure S5.**
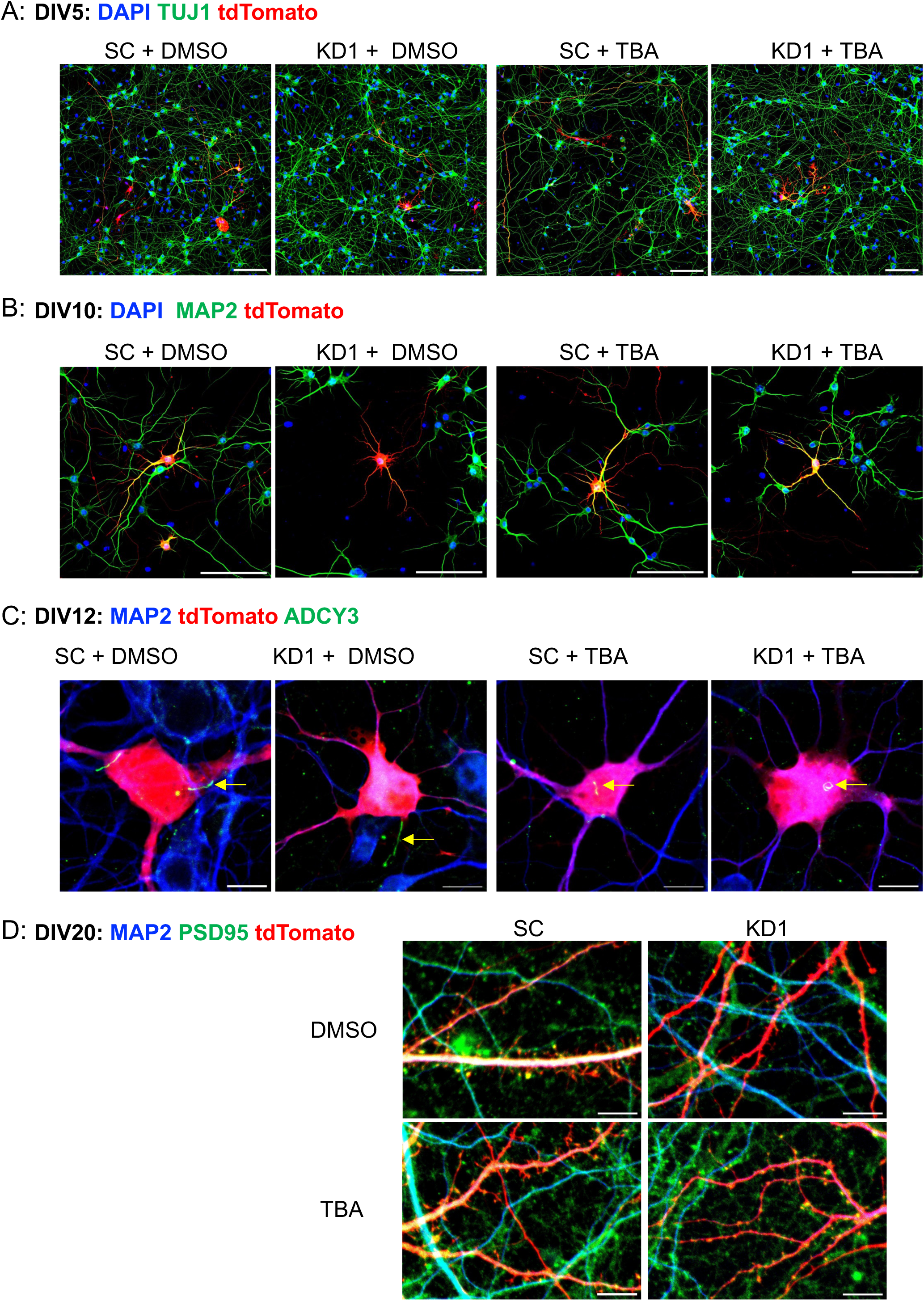
HDAC6 inhibition rescues cilium and spine defects but not neurite defects in LUZP1-deficient neurons. Related to Figure 4. Yellow arrowheads indicate ACDY3^+^ cilia. Scale bars, 100 μm (A and B) and 10 μm (C and D).

**Figure S6.**
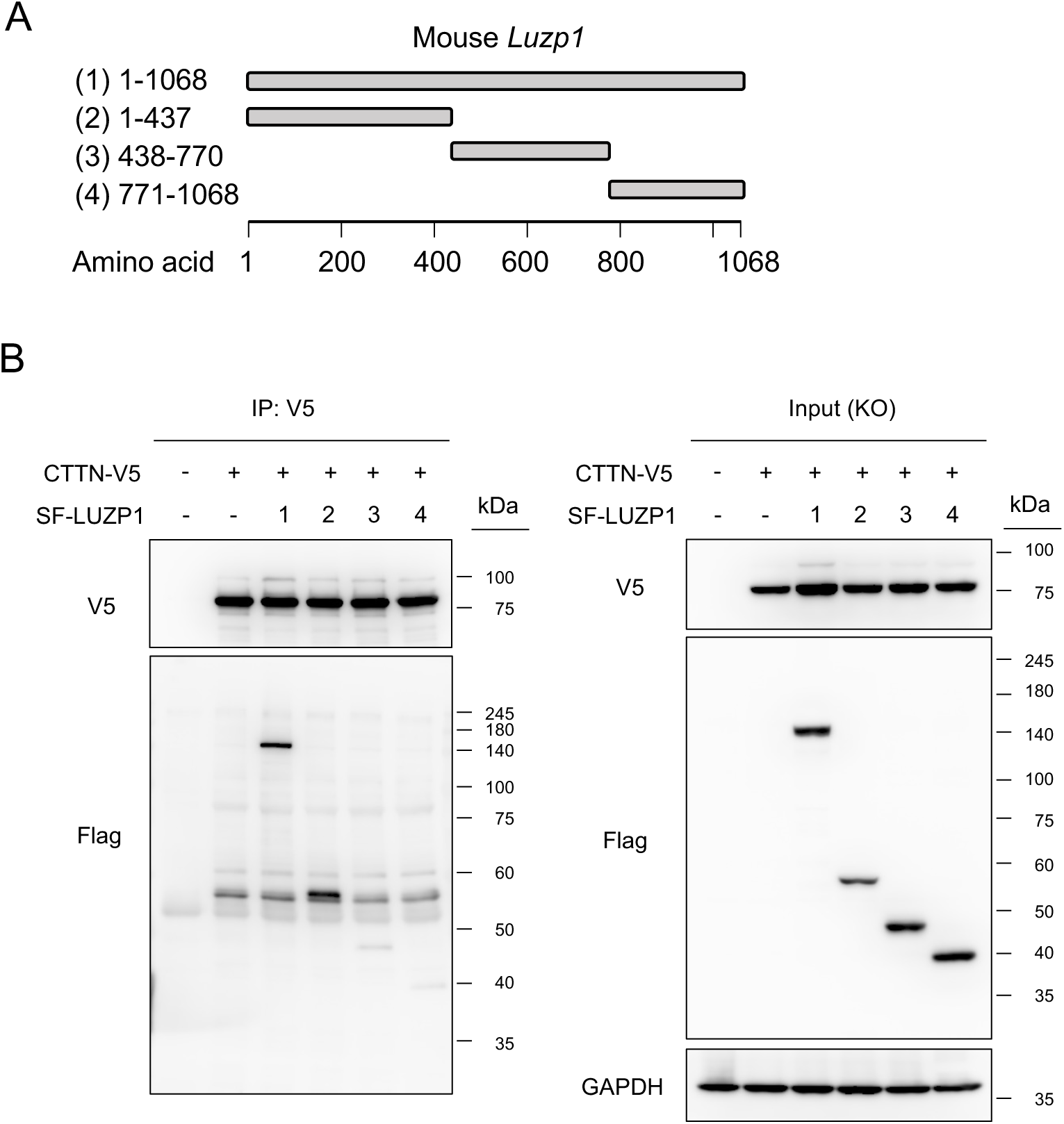
The N-terminal region of LUZP1 is required for interaction with CTTN. Related to Figure 5. (A) Schematic of WT and truncated LUZP1 constructs. (B) Immunoprecipitation assay testing interaction of CTTN with WT or truncated LUZP1 in LUZP1-KO HEK293T cells. Two independent experiments gave similar results.

**Figure S7.**
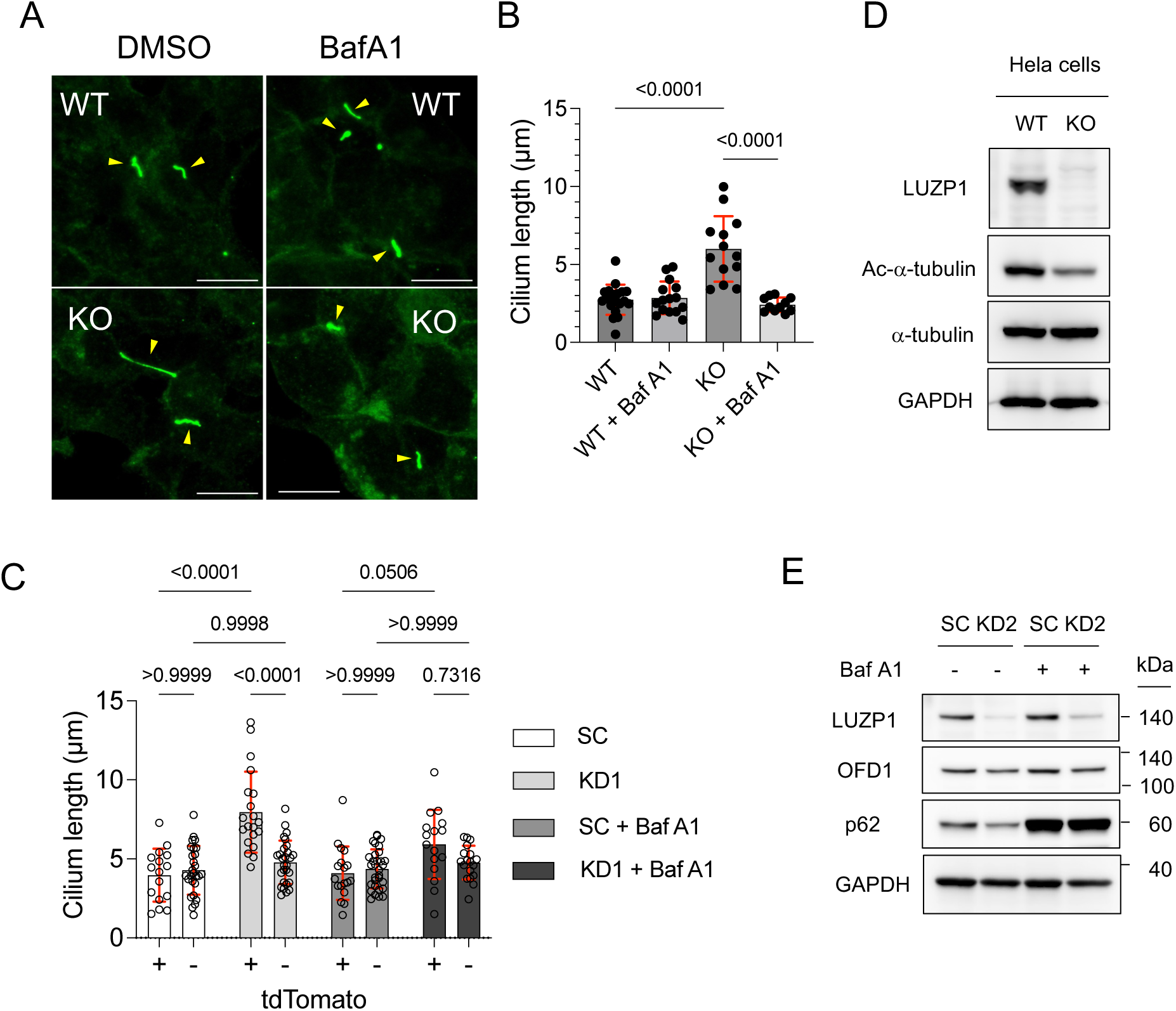
LUZP1 knockdown promotes ciliary elongation through autophagy. Related to Figure 7. (A and B) Arl13b^+^ cilia images and cilium length quantification in WT and LUZP1-KO HEK293T cells. (C) Cilium length in WT and LUZP1-knockdown hippocampal neurons with or without Baf A1. KD1 shRNA vectors were transfected into hippocampal neurons at DIV9, followed by Baf A1 treatment for 6 hours at DIV12 before fixation. In SC-knockdown groups, tdTomato-negative neurons showed no significant change in cilium length compared with tdTomato-positive neurons. (D) Validation of CRISPR-Cas9-mediated LUZP1 knockout in HeLa cells. (E) OFD1 and p62 immunoblots in control and LUZP1-knockdown hippocampal neurons. Hippocampal neurons were infected with AAV9-U6-shRNA virus carrying scramble or KD2 sequences at DIV9 and harvested at DIV18. Neurons were also treated with Baf A1 for 6 hours before harvest. Scale bars: 10 µm. Data are presented as means ± SD. Two-tailed unpaired t-tests (B). Exact P values are shown.

**Figure S8.**
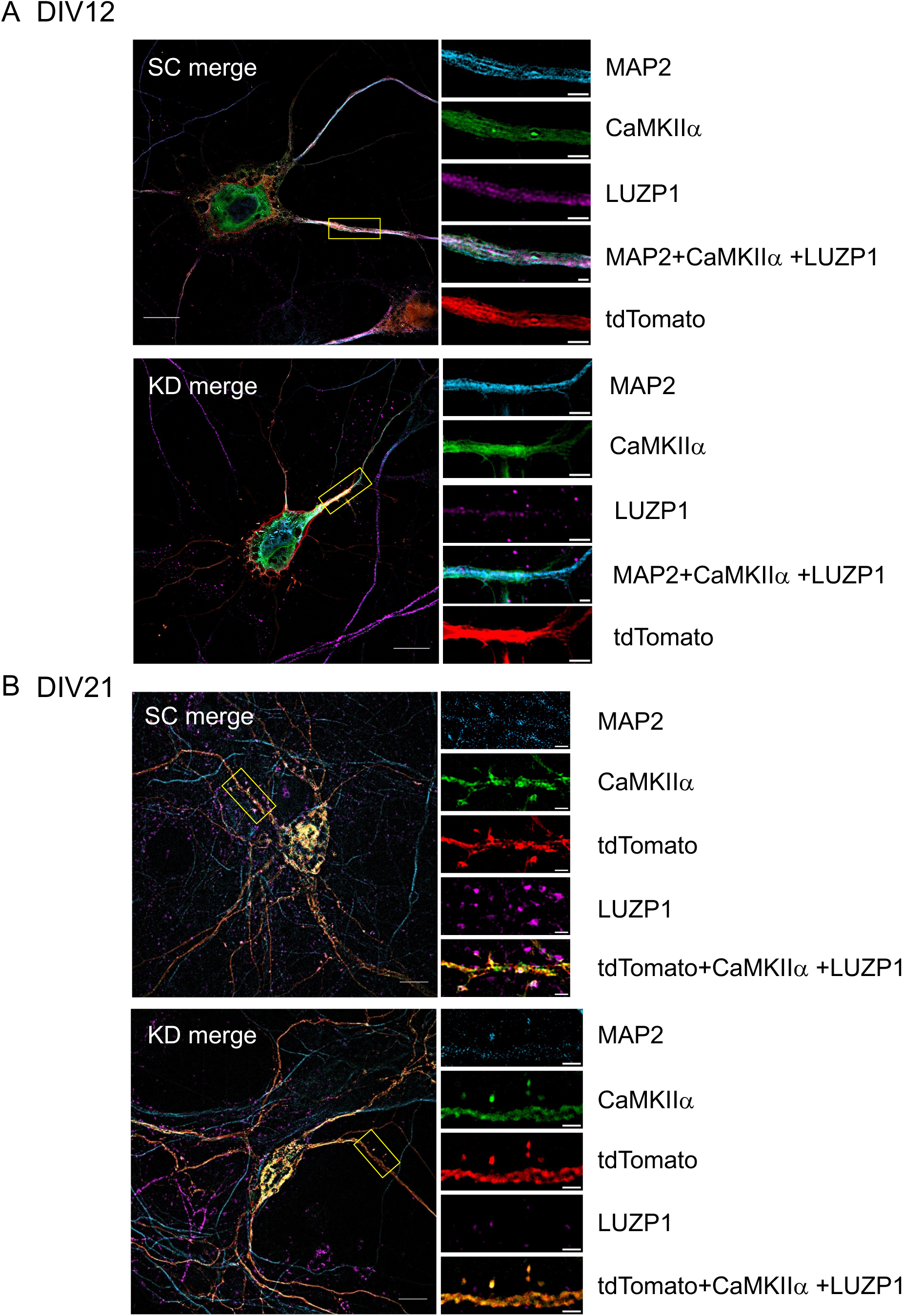
LUZP1 knockdown alters the CaMKIIα subcellular localization in dendrites. Related to Figure 8. (A and B) CaMKIIα subcellular localization in WT and LUZP1-knockdown hippocampal neurons. Neurons were transfected with KD1 shRNA vectors at DIV9 or DIV18 and fixed at DIV12 or DIV21, respectively. CaMKIIα subcellular localization was imaged using a ZEISS Lattice SIM5 super-resolution microscope.

